# Fly-casting with ligand–sliding and orientational selection to support the complex formation of a GPCR and a middle-sized flexible molecule

**DOI:** 10.1101/2022.02.28.482421

**Authors:** Junichi Higo, Kota Kasahara, Gert-Jan Bekker, Benson Ma, Shun Sakuraba, Shinji Iida, Narutoshi Kamiya, Ikuo Fukuda, Hidetoshi Kono, Yoshifumi Fukunishi, Haruki Nakamura

## Abstract

To elucidate computationally a binding mechanism of a middle-sized flexible molecule, bosentan, to a GPCR protein, human endothelin receptor type B (hETB), a GA-guided multidimensional virtual-system coupled molecular dynamics (GA-mD-VcMD) simulation was performed. This method is one of generalized ensemble methods and produces a free-energy landscape of the ligand-receptor binding by searching large-scale motions accompanied with stably keeping the fragile cell-membrane structure. All molecular components (bosentan, hETB, membrane, and solvent) were represented with an all-atom model, and sampling was carried out from conformations where bosentan was distant from the binding site in the hETB’s binding pocket. The deepest basin in the resultant free-energy landscape was assigned to the native-like complex conformation. The obtained binding mechanism is as follows. First, bosentan fluctuating randomly in solution is captured by a tip region of the flexible N-terminal tail of hETB via nonspecific attractive interactions (fly-casting). Bosentan then occasionally slides from the tip to root of the N-terminal tail (ligand–sliding). In this sliding, bosentan passes the gate of the binding pocket from outside to inside of the pocket with accompanying a quick reduction of the molecular orientational variety of bosentan (orientational selection). Last, in the pocket, ligand–receptor attractive native contacts are formed, and eventually the native-like complex is completed. The bosentan-captured conformations by the tip- and root-regions of the N-terminal tail correspond to two basins in the free-energy landscape, and the ligand–sliding corresponds to overcoming a free-energy barrier between the basins.

## Introduction

G protein-coupled receptors (GPCRs) are membrane proteins constructing a large evolutionarily related protein family with a variety of molecular functions^1^. In general, GPCR has seven transmembrane helices (TMH1 to TMH7), which are embedded in membrane. GPCRs activate cellular responses via detecting molecules outside the cell. Because GPCRs are related to many diseases, they have been important targets for drugs^2^. Endothelin-1 (ET1) is a peptide (21 residues long) with a strong vasoconstrictor action discovered in humans^3^. For exerting the activity, ET1 transmits signals by interacting with two GPCRs: The endothelin type A (ETA)^4^ and endothelin type B (ETB)^5^ receptors. The complex structure of human endothelin type-B receptor (hETB) and ET1 was solved using X-ray crystallography^6^. In this complex, ET1 is bound to a binding pocket of hETB on cytoplasmic side.

Bosentan competes with ET1 when binding to human ETA (hETA) and human hETB (hETB), and inhibits the ET1’s vasoconstriction effect as an antagonist^7,8,9^. The tertiary structure of the bosentan-hETB complex was solved by X-ray crystallography^10^. In this structure, bosentan binds to the binding pocket of hETB more deeply than ET1. However, the interaction patterns between the C-terminal segment of ET1 and aminoacid residues in the binding pocket of hETB are similar with those in the bosentan–hETB complex.

Bosentan is a medium-sized (551.6 Da) drug molecule and relatively large comparing to other commercial drug molecules. This molecule is flexible because it has long flexible sidechains (figure 1**a**) whereas the central ring, to which the sidechains are connected, is stiff. Thus, bosentan will adopt temporal complex conformations (encounter complexes) before reaching the deep position of the hETB’s pocket (i.e., the native-complex position). To investigate those temporal conformations, a special molecular dynamics (MD) simulation is suitable that can sample various conformations in the complex-formation process.

**Fig. 1.**
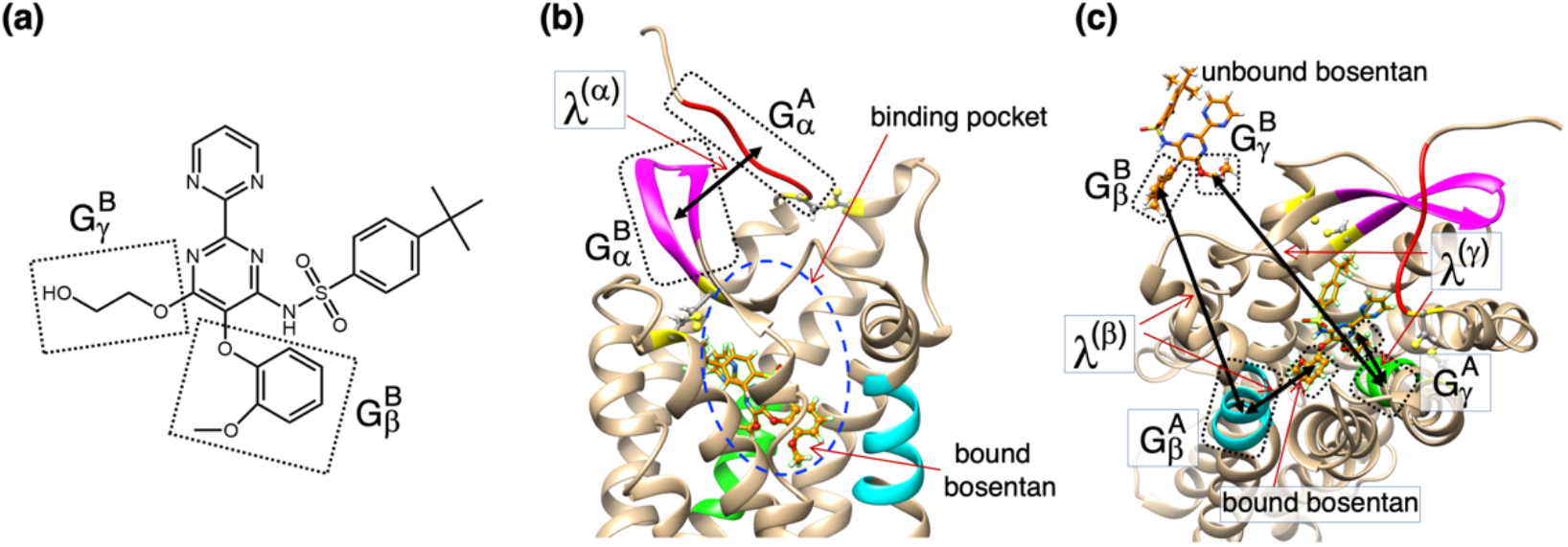
(A) Chemical structure of bosentan. Two atom groups 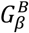 and 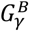, used to set reaction coordinates (RCs) *λ*^(*β*)^ and *λ*^(*γ*)^, are shown. (B) The first RC, *λ*^(*α*)^, defined by distance between centroids of atom groups 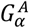 (red-colored segment in the N-terminal tail of hETB surrounded by dotted line; residues 85–89) and 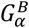 (magenta-colored segment in *β*-hairpin of hETB surrounded by another dotted line; residues 243–254). Binding pocket of hETB is indicated by blue-colored broken-line circle. Bound bosentan is shown at the bottom of the pocket. Green and cyan regions are defined in panel (C). (C) The second and third RCs, *λ*^(*β*)^ and *λ*^(*γ*)^, respectively. *λ*^(*β*)^ is defined by distance between centroids of 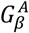 (a cyan-colored part of the fifth transmembrane helix TMH5; residues 273–281) and 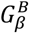 (a part of bosentan surrounded by dotted line). *λ*^(*γ*)^ is defined by 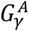 (a green-colored part of the seventh transmembrane helix TMH7; residues 372–379) and 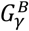 (other part of bosentan surrounded by the other dotted line). Bound and unbound bosentans are displayed.

Recently, we introduced a generalized ensemble method, a multi-dimensional virtual-system coupled molecular dynamics (mD-VcMD) simulation^11^, which enhances conformational sampling with repetition of simulations (iteration) in a multi-dimensional (mD) reaction-coordinate (RC) space. Repeating the iterations, the sampled volume in the mD-RC space is expanded and sampled conformations (snapshots) are collected from the iterations. Importantly, mD-VcMD assigns a statistical weight (thermodynamic weight equilibrated at a simulation temperature) to each of the snapshots. Therefore, the ensemble of snapshots is equilibrated, and any equilibrated physicochemical quantity is computable from the ensemble if the quantity is expressed by the system’s coordinates. We call this simulation method the “original mD-VcMD method” or simply the “original method” in this paper. The original method was applied to a large and complicated system consisting of ET1 and hETB, where hETB was embedded in an explicit membrane and the membrane was immersed in an explicit solvent^12^.

Although the original mD-VcMD was applicable to the large and complicated system, we encountered a difficulty: Once a poorly sampled region emerged in the mD-RC space in an iterative simulation, the conformation might be trapped in this region in the subsequent iteration. Thus, a more robust method was required to proceed the sampling. To overcome this difficulty, we introduced a GA-guided multi-dimensional virtual-system coupled molecular dynamics method (GA-mD-VcMD)^13^, which is an extension of the original mD-VcMD, where “GA” is an abbreviation of genetic algorithm. We applied this method to some biological systems, which consist of a protein and ligand in an explicit solvent^14,15,16^.

From the original mD-VcMD applied to binding of ET1 to hETB, two binding mechanisms were found: Fly casting^17,18,19^ and conformational selection^20^. The receptor hETB has an intrinsically disordered N-terminal tail. The floating ET1 in solution was captured by the fluctuating N-terminal tail (fly-casting mechanism) because the interactions between ET1 and the hETB’s N-terminal were attractive^12^. We note that the root of the N-terminal tail is located at the gate of the binding pocket of hETB. This indicates that once ET1 is captured by the N-terminal tail, the ligand is restricted in a narrow volume around the gate of the binding pocket, resulting in increase of the possibility that ET1 contacts the gate of the binding pocket. Next, only a fraction of various orientations of ET1 were allowed to enter the hETB pocket (conformational selection). This suggests that the pocket size is not large enough for ET1 to rearrange the molecular orientation in the pocket, and that only ET1 with molecular orientations advantageous for the native complex formation is accepted.

In the present study, we sampled the interactions between the medium-sized molecule bosentan and hETB by the GA-mD-VcMD simulation. Except for the difference of ligand and simulation method, there are two differences between the previous and present studies. (i) We prolonged the intrinsically disordered N-terminal tail of hETB of the X-ray structure^6,10^: Five and ten residues long for the previous and the current studies, respectively^12^. This tail extension varies the feature of the fly-casting mechanism as shown later. Because GPCRs have a long N-terminal tail in general^21^, this extension of the tail may support experimental study on the role of the N-terminal tail of GPCRs^22,23^. Second, the initial simulation conformations for the present study were those where bosentan was distant from hETB while those for the previous study were the native-like complex conformations of ET1 and hETB. In general, the difference of initial conformations significantly influences the resultant conformational ensemble. The current study is not guaranteed if the native-like conformations are sampled. The nativelike complex should be *discovered* as the lowest free-energy conformation out of many possible ones. Contrarily, in the previous study, the native-like complex is always involved in the resultant ensemble, which means that the native-like complex may not necessarily correspond to the lowest free-energy conformation because a lower free-energy conformation may be overlooked in the sampling.

Our GA-mD-VcMD simulation produced a free-energy landscape of the bosentan–hETB system, and probability distribution functions of some quantities were calculated from the resultant conformational ensemble. Because the free-energy landscape covered both unbound conformations and the native-like complex structures, and because the lowest free-energy conformation corresponded to the native-like complex structure, we could analyze entirely the complex formation. We discuss the generality of the binding mechanism found in bosentan-hETB to the ligand–GPCR binding. Because bonentan is a drug molecule, the present study helps drug development research not only by proposing the lowest free-energy conformations but also by investigating the binding process.

## Methods

The aim of this work is to investigate the complex-formation process of bosentan and hETB by the GA-mD-VcMD method. First, we briefly explain this method, which provides a conformational ensemble of the system and a thermodynamic weight at the simulation temperature *T* (300 K in the present study), which is assigned to conformations stored in the ensemble. Details of GA-mD-VcMD have been explained elsewhere^13,14,15,16^. Next, we explain the molecular system which consists of hETB, bosentan, membrane (cholesterol and POPC lipid molecules), and solvent (water molecules and ions). Then, we describe multiple reaction coordinates introduced and the initial conformations for simulation. Lastly, we account for the method to calculate the spatial density of bosentan around hETB.

### GA-mD-VcMD and a conformational ensemble

To sample extensively both the unbound and bound conformations of the system, GA-mD-VcMD controls the system’s motions in a multi-dimensional (mD) space constructed by multiple reaction coordinates (RCs). Definition of an RC is presented in Section 1 of Supplementary Information (SI) and figure S1. We introduce three RCs denoted as *λ*^(*h*)^ (*h* = *α*,*β*, or *γ*), whereas the dimensionality of the RC space is arbitrary in theory^13^. Therefore, the word “multi-dimensional (mD)” indicates three-dimensional (3D) in the present study. The actual definition for the three RCs is given later.

Conformational sampling of the system by GA-mD-VcMD at 300 K produces a distribution (density) of the system in the mD-RC space (not in the real space). Because the conformational space of a biological molecular system is wide, the GA-mD-VcMD simulation is executed iteratively, by which the sampled volume of 3D-RC space is expanded. Here, the density at a 3D-RC position [*λ*^(*α*)^,*λ*^(*β*)^,*λ*^(*γ*)^] is denoted as *Q_cano_*(*λ*^(*α*)^,*λ*^(*β*)^,*λ*^(*γ*)^), and the density computed from the *M*-th iteration may be expressed as 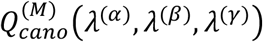 when necessary.

The simulation is terminated when a convergence criterion is satisfied for *Q_cano_*(*λ*^(*α*)^,*λ*^(*β*)^,*λ*^(*γ*)^). Snapshots of the system are saved from all iterations. After the convergence, a thermodynamic weight at the simulation temperature *T* is assigned to each saved snapshot based on *Q_cano_*(*λ*^(*α*)^,*λ*^(*β*)^,*λ*^(*γ*)^). Therefore, the snapshots construct a thermally equilibrated conformational ensemble (canonical ensemble) at *T* (300 K).

### Molecular system studied

We generated the molecular system consisting of bosentan, hETB, cholesterols surrounding hETB, POPC bilayer where hETB is embedded, and an explicit solvent (water molecules, *Na*^+^ and *Cl*^-^ ions). All of them were packed in a periodic boundary box. Details for the system preparation are given in Section 2 of SI. figure S2 is a diagram outlining the generation of the molecular system.

Two reference crystal structures are available for generating the molecular system for the simulation: The bosentan–hETB (PDB ID 5xpr; 3.6 Å resolution) and K8794–hETB (5×93; 2.2 Å) complex structures. Importantly, the two crystal structures are considerably similar each other (figure S3**a**) because K8794 is an analog of bosentan (figure S3**b**). Because K8794–hETB complex has a better resolution than bosentan–hETB complex, we used the K8794–hETB complex to prepare for the molecular system.

First, we replaced K8794 by bosentan. This complex conformation is referred to “Complex structure of bosentan–hETB” in figure S2. In the crystallography, some mutations were applied to the amino-acid sequence of hETB to increase the quality of the diffraction data^10^. Then, we returned the hETB’s sequence to the wild type, and embedded the complex in the POPC bilayer. Four cholesterols were introduced into the hETB– membrane interface (see Section 2 of SI). Then, the system was immersed in a periodic boundary box filled by water molecules. The x-, y-, and z-coordinate axes for specifying the atomic positions were set parallel to the sides of the periodic box (the x-y plain was parallel to the membrane surface). The box dimension was 71.33 Å (x-axis), 71.33 Å (y-axis), and 132.17 Å (z-axis). Finally, sodium and chlorine ions were introduced by replacing water molecules randomly with the ions to neutralize the system at a physiological ion concentration of 130 mM. This conformation was named “Complex structure in periodic box” in figure S2. The total number of atoms of the system was 69,062: numbers of protein and bosentan atoms were 5,289 and 68, respectively; four cholesterols (296 atoms); 127 POPC molecules (17,018 atoms); 15,440 water molecules (46,320 atoms); and 28 sodium and 43 chlorine ions.

After energy minimization of the above system, followed by NVT and NPT simulations, we obtained the complex structure shown in figure S4, whose resultant periodic-box size was 69.37 Å × 69.37 Å × 140.29 Å. This is represented as “Complex structure in figure S4 (native-complex structure)” in figure S2. Because this complex structure is a relaxed conformation of the native complex, bosentan was located at a position close to that in the X-ray structures (PDB ID: 5xpr and 5×93). Therefore, we call this conformation as the “native complex structure” in the present study. It should be noted that this native complex structure is *not* the initial conformation of the current GA-mD-VcMD simulations. As explained later, we generated conformations where bosentan was completely separated from the binding pocket of hETB. They were the conformations that we used as the initial simulation conformations of GA-mD-VcMD.

### Set of Three RCs Introduced to Control System’s Motions

Before describing the generation of the initial conformations of GA-mD-VcMD, we explain concretely the set of three RCs (*λ*^(*h*)^; *h* ∈ {*α*,*β*,*γ*}) introduced for the current study. Setup of RCs is essential to increase the sampling efficiency though RCs can be set arbitrary in theory. We imposed three conditions on the RCs: The RCs can control (i) the gate opening/closing of the hETB’s binding pocket, (ii) the ligand approaching/departing to the receptor, and (iii) the ligand orientation relative to the receptor.

Although various RCs are possible to satisfy the three conditions above, we set them in a straightforward manner as shown in figures 1**b** and **c**, and the exact specification for the atom groups are given in the figure caption. We briefly explain the roles of the RCs. For the condition (i), two atom groups 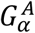 and 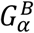 (figure 1**b**) are used to define the first RC *λ*^(*α*)^ that describes open /close of the gate of the hETB’s binding pocket. figure 1**c** shows atom groups 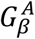 and 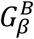 to define *λ*^(*β*)^, as well as 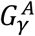 and 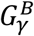 to define *λ*^(*γ*)^. The groups 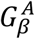 and 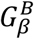 are close to each other in the native complex, and 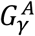 and 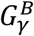 are as well. Contrarily, those atom groups are distant from each other in an unbound conformation. Therefore, the variations of *λ*^(*β*)^ and *λ*^(*γ*)^ control bosentan approaching/departing to hETB: The condition (ii) is satisfied. Lastly, the condition (iii) is satisfied when *λ*^(*β*)^ increases/decreases with decreasing/increasing *λ*^(*γ*)^. Figure 1**a** is closeup of bosentan to clarify 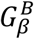 and 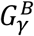.

### Initial conformations for GA-guided mD-VcMD

We explain generation of the initial conformations of GA-mD-VcMD. Our aim is to explore the conformational space from the unbound to bound states. Thus, the initial conformations should not be the native complex structure but be a completely unbound conformations. Since we had only the native complex structure (figure S4) at this stage, we needed to generate the unbound conformations from the complex structure. For this purpose, we performed ten simulations (called “separation simulations”) starting from the native complex structure with imposing three requirements to RCs to the system’s conformation: 24.0 Å ≤ *λ*^(*α*)^ ≤ 25.0 Å, 60.0 Å ≤ *λ*^(*β*)^ ≤ 65.0 Å, and 60.0 Å ≤ *λ*^(*γ*)^ ≤ 65.0 Å. The three RCs for the native complex structure are *λ*^(*α*)^ = 8.22 Å, *λ*^(*β*)^ = 8.35 Å, and *λ*^(*γ*)^ = 6.21 Å, respectively. During the separation simulations, bosentan moved from the binding pocket to unbound conformations. The resultant conformations are shown in figure S5, where bosentans are completely separated from hETB, which are represented “Unbound conformations in figure S5” in figure S2.

Next, for further relaxing those unbound conformations, we performed 2,000 runs in total: 200 (1 × 10^6^ steps at 300 K) runs starting from each of those ten unbound conformations with different random seeds to the initial atomic velocities. In the 2,000 runs, we applied the following three restrictions to RCs: *λ*^(*α*)^ ≤ 25.0 Å, *λ*^(*β*)^ ≤ 65.0 Å, and *λ*^(*γ*)^ ≤ 65.0 Å. The upper limits were introduced to prevent bosentan from flying further away from hETB in the periodic box. We refer to those 2,000 runs as “diffusion simulations”. The last snapshots from 2,000 runs were used as the initial conformations of the GA-mD-VcMD simulations. Figure S6 plots the positions of the last snapshots of the simulations in the 3D-RC space and the native complex structure in the 3D-RC space. Those conformations are represented as “Distributed conformations in figure S6” in figure S2.

It should be noted that figure S6 demonstrates that the conformations sampled in the diffusion simulations did not reach the native complex structure. We emphasize that all the current results were not influenced by the structural features of the native complex because the 2,000 initial conformations are those diffused freely from the ten completely unbound conformations (figure S5).

### GA-mD-VcMD simulations

As mentioned in the INTRODUCTION–section, GA-mD-VcMD enhances conformational sampling in the 3D-RC space, and the thermodynamic weights of sampled conformations at the simulation temperature (300 K) are computed using the converged distribution *Q_cano_*(*λ*^(*α*)^,*λ*^(*β*)^,*λ*^(*γ*)^).

In the original mD-VcMD^12^ and GA-mD-VcMD^13^, each RC axis *λ*^(*h*)^ (*h* ∈ {*α*,*β*, *γ*}) is divided into small 1D zones (the zone was named originally as “virtual zone”^13^), and accordingly the 3D-RC space is divided into small 3D zones. Table S1 lists the actual division of each 1D axis into zones. During a short time–interval Δ*τ_trns_* of simulation, the system’s conformation freely moves only in a single 3D zone with a restoring force outside the zone. Then, an inter-zone transition from the zone to one of neighboring zones is attempted every time interval. In the present study, we set Δ*τ_trns_* = 20 ps, which corresponds to 1 × 10^4^ simulation steps with time step of 2 /s. The interzone transition probabilities are determined using *Q_cano_* ^12,13^. In this way, the conformation moves in a wide 3D-RC space by the inter-zone transitions.

We performed *N_run_* (= 2,000) runs at 300 K in parallel starting from different conformations in each iteration to further increase the sampling efficiency. A simple integration of the *N_run_* trajectories can be regarded as a single, long trajectory by connecting the trajectories in arbitrary order^24,25^. The *N_run_* initial conformations for the first iteration of GA-mD-vcMD are the last conformations from the diffusion simulations (figure S6). We obtain snapshots from the first to *M*-th iterations when the *M*-th iteration has finished. The *N_run_* initial conformations for the (*M* + 1)-th iteration are selected from those snapshots so that they distribute as evenly as possible in the 3D-RC space. On the other hand, when a certain RC region is poorly sampled, we pick more snapshots from or near the poorly sampled RC region than from well-sampled RC regions^13^.

We set the simulation length of each run to 1 × 10^6^ steps (1 × 10^6^ × 2 fs = 2.0 ns). Thus, the total length for an iteration is 4.0 μs (=2.0 ns × 2,000). We performed 55 iterations in the present study, which corresponds to 220.0 μs (4.0 μs × 55) in total. A snapshot was stored every 1 × 10^5^ steps (0.2 ns) in each run, and then 1.1 × 10^6^ (220.0 μs/0.2 ns) snapshots were stored in total. We calculated distribution functions of various quantities at the simulation temperature (300 K) from the resultant ensemble of snapshots.

The GA-mD-VcMD algorithm^13^, which was initially available in a MD simulation program omegagene/myPresto^26^, has now been implemented in GROMACS^27^ using the PLUMED plug-in for the current study^28^. The simulation condition is as follows: LINCS^29^ to constrain the covalent-bond lengths related to hydrogen atoms, the Nosé–Hoover thermostat^30,31^ to control the simulation temperature, the zero-dipole summation method^32,33,34^ to compute the long-range electrostatic interactions accurately and quickly, a time-step of 2 fs (Δ*t* = 2 fs), and simulation temperature of 300 K. All simulations were performed on the TSUBAME3.0 supercomputer at the Tokyo Institute of Technology with using GP-GPU.

The force fields used are AMBER99SB-ILDN^35^ for hETB and AMBER LIPID14^36^ for the cholesterol and POPC lipid molecules, TIP3P^37^ for water molecules and Joung–Cheatham model^38^ for ions (chloride and sodium), respectively. The bosentan’s force field was prepared as follows: First, the bosentan’s atomic partial charges were calculated quantum-chemically using Gaussian09^39^ at the HF/6-31G* level, followed by RESP fitting^40^. Then, the obtained partial charges were incorporated into a GAFF (general amber force field) force-field file^41^, which was designed to be compatible with the conventional AMBER force-fields. The chemical structure of bosentan is shown in figure 1A.

### Spatial density of bosentan around hETB

GA-mD-VcMD simulation produces a canonical ensemble of conformations each of which a thermodynamic weight at 300 K is assigned to. Therefore, using the ensemble, we can calculate a spatial density of bosentan’s centroid around hETB at 300 K using the ensemble. The detailed method to calculate the spatial density is presented in Subsection 3.1 of SI.

We explain briefly notations used to express the spatial density. The 3D real space (not the 3D-RC space) is divided into cubes (size is 3 Å × 3 Å × 3 Å), whose centers are denoted as ***r**_cube_*. The spatial density *ρ_CMb_*(***r**_cube_*) is the probability of detecting the bosentan’s centroid ***r**_CMb_* in the cube centered at ***r**_cube_*.

## Results and Discussion

We repeated 55 iterations of GA-mD-VcMD with saving 1.1 × 10^6^ snapshots from the first to 55-th iterations in total (see Methods section). First, we show that the GA-mD-VcMD produced a distribution of the system that covered both the unbound and nativelike complex conformations. We next show that the highest density spot (the lowest free-energy spot) corresponds to the native-like complex structure. We then analyzed the binding process from the unbound to intermediates, and from the intermediates to the native-like complex. For this purpose, the binding process is divided into three stages: One is the early stage of binding in that bosentan is outside the binding pocket. The second is the stage where bosentan is passing the gate of the binding pocket. The last is the stage where bosentan is moving in the binding pocket.

### Conformational distribution in 3D-RC space

GA-mD-VcMD produced the distribution *Q_cano_*(*λ*^(*α*)^,*λ*^(*β*)^,*λ*^(*γ*)^) equilibrated at 300 *K* in the 3D-RC space (figure S8). Importantly, the distribution covered both the unbound and bound conformations, and the native-complex structure was located at the periphery of the lowest free-energy basin. Section 4 of SI explains the features of this distribution.

The method to check the convergence of the distribution is presented in Section 5 of SI. Figure S9 shows that iterations 1–10 (figures S9**a**-**c**) sampled a large volume of the 3D-RC space involving the unbound and bound conformations. Therefore, the simulation could be quitted at the tenth iteration. However, to obtain snapshots sufficient for analysis, we continued the simulation up to the 55-th iteration. Remember that we stored a snapshot every 0.2 ns (100,000 steps) of simulation.

Next, we compared the distribution *Q_cano_*(*λ*^(*α*)^,*λ*^(*β*)^,*λ*^(*γ*)^) with that obtained from the previous study of the ET1–hETB system^12^. See Section 6 of SI for details of the comparison method. Figure S10 shows that the free-energy slope along the *λ*^(*α*)^ axis (the motion along the gate opening/closing motion of the hETB’s binding pocket) is gentler than those along the *λ*^(*β*)^ or *λ*^(*γ*)^ axis (the bosentan’s approaching/departing motions to hETB). This result agrees with that from the previous study. However, the slope in the current bosentan–hETB landscape was about one-tenth of that in the previous ET1-hETB landscape. We discuss later this inconsistency between the two studies.

### Three-dimensional (3D) spatial distribution of bosentan around hETB

Whereas GA-mD-VcMD samples the mD-RC space, the conformational distribution in the mD-RC space does not always provide a concrete or intuitive image for the molecular conformation of the system. Therefore, we converted the distribution *Q_cano_*(*λ*^(*α*)^,*λ*^(*β*)^,*λ*^(*γ*)^) (figure S8) to the conformational distribution *ρ_CMb_*(***r**_cube_*) defined in the 3D real space using a thermodynamic weight assigned to all the sampled snapshots using equation 31 of Ref. 13. See Subsection 3.1 of SI for the method to calculate *ρ_CMb_*(***r**_cube_*). Figure 2 illustrates the spatial patterns of *ρ_CMb_*(***r**_cube_*) of the bosentan’s centroid ***r**_CMb_* around hETB. Importantly, the highest-density spot (the red-colored contours; *ρ_CMb_*(***r**_cube_*) ≥ 0.5) well agreed with the position of bosentan’s centroid in the native complex structure. This result is important because the reliability of the simulation data reduces unless the highest-density spot is assigned to the native bosentan’s position.

**Fig. 2.**
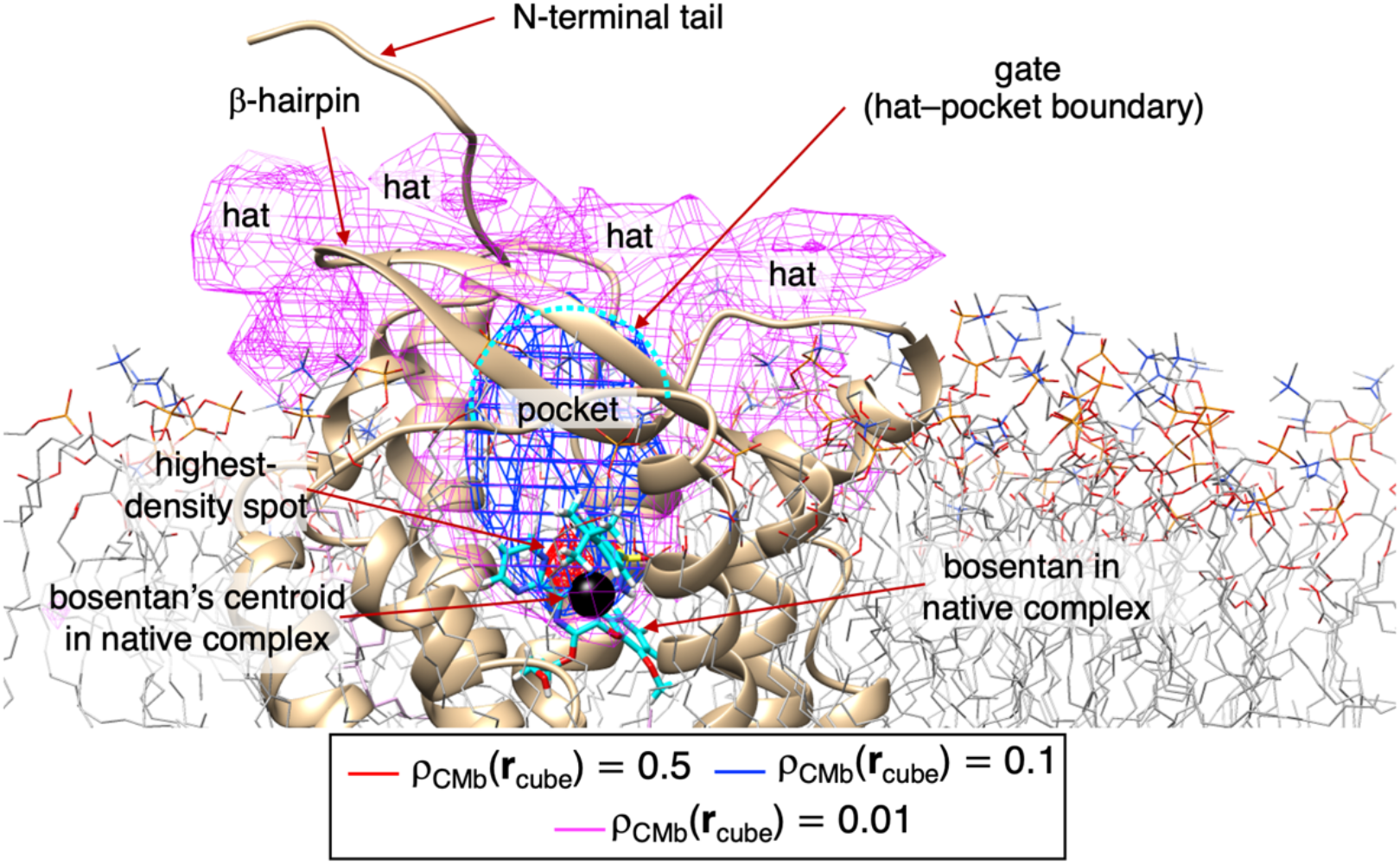
Spatial density, *ρ_CMb_*(***r**_cube_*), of bosentan’s centroid around hETB. Iso-density map is presented at three contour levels by different colors as shown in inset. Molecular structure is the native complex (figure S4) omitting solvent. Bosentan is shown by cyan-colored stick model, and its centroid is shown by small black sphere. N-terminal tail and *β*-hairpin of hETB are indicated with labels. Red-colored contours are named “highest-density spot”. Blue-colored ones (*ρ_CMb_*(***r**_cube_*) ≥ 0.1) specify well the binding pocket of hETB. Magenta-colored region above the pocket was named “hat”. Cyan-colored broken-line indicates boundary between hat–pocket boundary, which is called “gate” of the binding pocket. See also figure S11, which illustrates *ρ_CMb_*(***r**_cube_*) with adding two low-density contours of *ρ_CMb_*(***r**_cube_*) ≥ 0.001 and 0.0001.

The binding pocket corresponds to the region of *ρ_CMb_*(***r**_cube_*) ≥ 0.1 (bluecolored contours), which involved the highest-density spot. Outside of the binding pocket was characterized by a lower density of *ρ_CMb_*(***r**_cube_*) ≥ 0.01 (magenta-colored ones), which was named “density hat” or “hat” simply. We denote the hat–pocket boundary (cyan-colored broken line in Figure 2) as “gate” of binding pocket.

Figure S11 illustrates the spatial density for the whole sampled region. Bosentan was distributed even in far regions from hETB as indicated by the low density: *ρ_CMb_*(***r**_cube_*) ≥ 0.0001. The bosentan’s density increased rapidly with approaching the binding pocket from the far region. This figure demonstrates that GA-mD-VcMD is powerful in sampling the wide conformational space efficiently.

### RMSD of Bosentan between Snapshot and the Native Complex

Above we showed visually agreement of the computed highest-density spot and the bosentan’s native-complex position. Here, to analyze quantitatively the agreement, we calculated two types of root-mean-square-deviation *RMSD* of bosentan between a snapshot and the native complex: *RMSD* = *RMSD_whole_* or *RMSD* = *RMSD_core_*. *RMSD_whole_* is calculated using all the heavy atoms in bosentan, and *RMSD_core_* is done using the heavy atoms in the bosentan’s core region. Definition of the core region is given in figure 3**a**.

**Fig. 3.**
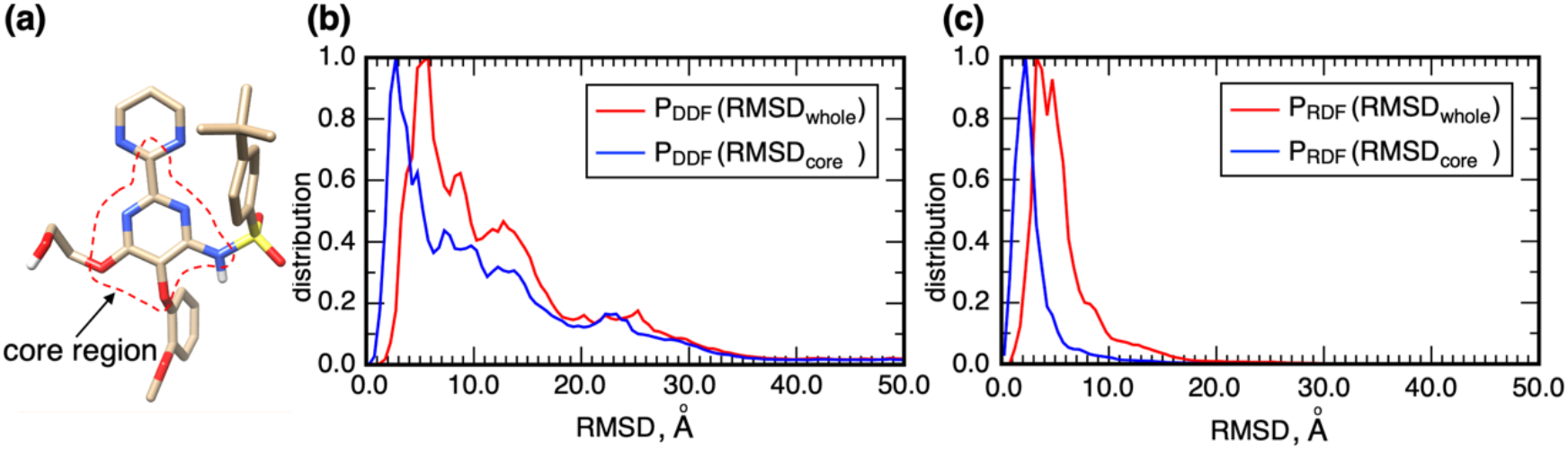
(**a**) Bosentan’s “core region”, which is surrounded by red-colored broken line. (**b**) Distance distribution function *P_DDF_*(*RMSD*) and (**c**) radial distribution function *P_RDF_*(*RMSD*) are presented for both of *RMSD_wbole_* and *RMSD_core_*. Definition of *RMSD*, *P_DDF_*, and *P_RDF_* is given in Subsection 3.2 of SI.

We calculated two probability distribution functions to analyze *RMSD*: Distance distribution function *P_DDF_*(*RMSD*) and radial distribution function *P_RDF_*(*RMSD*). See Subsection 3.2 of SI for details of *P_DDF_* and *P_RDF_*. The resultant distributions are presented in figure 3. For the whole bosentan, the highest-peak positions of *P_RDF_* and *P_DDF_* are at *RMSD_whole_* ≈ 3.25 Å and *RMSD_whole_* ≈ 5.75 Å, respectively. Similarly, for the core region, those are at *RMSD_core_* ≈ 2.25 Å and *RMSD_core_* ≈ 2.75 Å, respectively. The peak positions for the core region were smaller than those for the whole bosentan. This is because of the disorder of the longest sidechain of bosentan, as shown later.

### Bosentan–tail contact ratio

The structure of the N-terminal tail is not determined in the x-ray structures (PDB IDs: 5×93 and 5xpr) because the tail is intrinsically disordered. We presume that the tail may affect the binding process because the root of the tail is located at the entrance of the hETB’s binding pocket. Some experiments report that an extracellular loop of GPCRs plays a role of lid to block the gate of the binding pocket^42,43^.

Figure S7**b** highlights the gate of the hETB’s binding pocket. This figure shows that a half of the gate, which is covered by the white solid-line rectangle, is closed by the contact of the N-terminal tail and the *β*-hairpin, and the other half of the gate, which is covered by the white broken-line rectangle, is open. The definition of the N-terminal tail and the *β*-hairpin is given in the caption of figure S7**b**. We can imagine readily from this figure that the gate situation can vary during the fluctuations of the tail. Below, we investigate the tail’s fluctuations and the bosentan–tail contact.

First, to visualize the fluctuations of the N-terminal tail, we calculated a spatial density function *ρ_CMNt_*(***r**_cube_*) for the tip of the N-terminal tail. We also calculated the spatial density function *ρ_CMβh_*(***r**_cube_*) for the tip of the *β*-hairpin because the *β*-hairpin is the contact partner of the N-terminal tail to open/close the gate of the binding pocket. Figure S12**a** demonstrates that the N-terminal tail fluctuated widely with varying its orientation, which is a natural property of the intrinsically disordered segment. In contrast, the *β*-hairpin fluctuation is highly limited in a narrow volume as shown in figure S12**b**. Therefore, the gate opening/closing is induced by the large motions of the N-terminal tail (not by the fluctuation of *β*-hairpin).

Next, to investigate the bosentan–tail contact, we introduced a “cube-based bosentan–tail contact ratio” denoted as 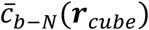. Remember that the bosentan’s centroid ***r**_CMb_* in a snapshot was assigned to a cube at ***r**_cube_*, which involved ***r**_CMb_*. Then 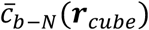 is the ratio that bosentan in ***r**_cube_* contacts to the N-terminal tail. The maximum and minimum of 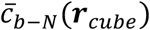 are 1.0 and 0.0, respectively. Detained procedure to calculate 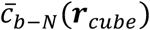 is given in Subsection 3.3 of SI.

Figure 4**a** illustrates the spatial patterns of 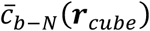 at three contour levels: 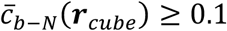, 0.7, and 0.9. The low-contact contours, 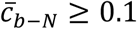, extended to regions distant from the gate of the binding pocket, which means that the N-terminal tail contacted bosentan even when bosentan was floating in solution. With bosentan approaching the pocket, the contact ratio increased quickly and reached 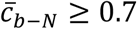. Because the region of 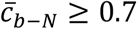 involved the density-hat region (*ρ_CMb_* ≥ 0.01; magenta-colored contour regions of figure 2), the contact may act as an attractive interaction. The high-contact region of 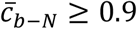 (magenta-colored contours in figure 4**a**) was located just above the pocket not in the hETB’s binding pocket. Interestingly, 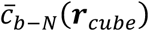 was considerably small in the pocket. The loss of the bosentan–tail contact in the pocket indicates that the role of the N-terminal tail is the recruit of bosentan into the pocket.

**Fig. 4.**
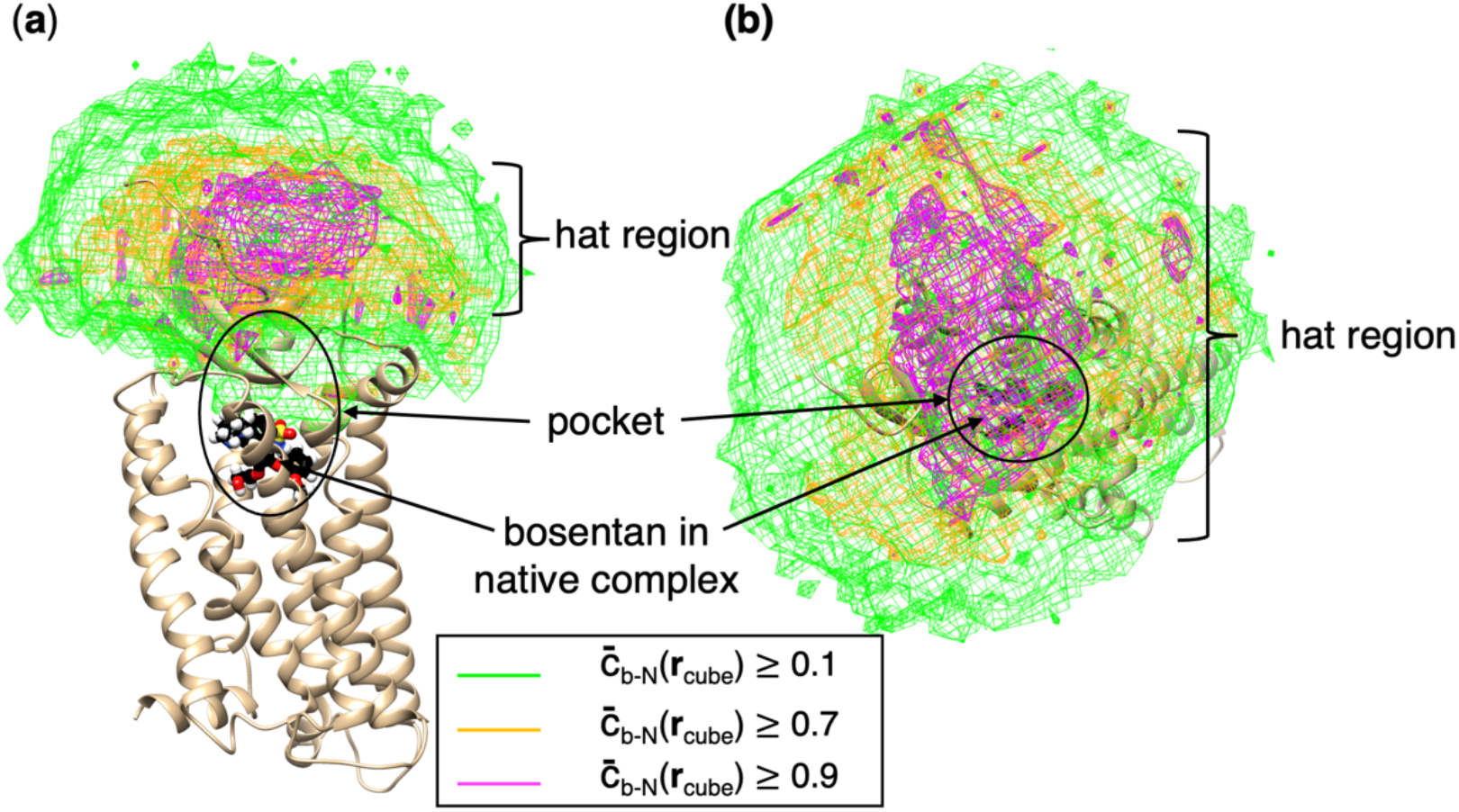
(**a**) Spatial patterns of cube-based bosentan–tail contact ratio 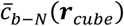 at three contour levels shown in inset. The “hat” region, “pocket”, and “bosentan in native complex” are also shown. (**b**) Spatial patterns viewed from above of the pocket. Shown structure is the native complex.

Remember that the gate-closing is brought by the contact of the N-terminal tail and the *β*-hairpin (figure S7**b**). In other words, the gate-opening is induced by the N-terminal tail fluctuating in solution away from the gate. Figure 4 indicate that the regions of 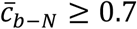 are widely extended far from the binding pocket. The gate-opening/closing, the large fluctuations of the N-terminal tail and the bosentan–tail contact formation are related mutually.

### Bosentan slides along the N-terminal tail upon binding

As shown in figure 4, the cube-based bosentan–tail contact ratio depended on the relative position of bosentan to hETB in the 3D real space. Here, we quantify the bosentan’s position relative to hETB by a quantity *r_bb_*, which is the distance between the bosentan’s centroid, ***r**_CMb,i_*, in snapshot *i* and that in the native complex 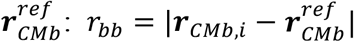. Figure S7**c** illustrates *r_bb_* in a snapshot. The smaller the *r_bb_*, the closer the bosentan to its natively binding position. Then, we divided the *r_bb_* axis into thirteen slices with thickness of 5 Å, which are denoted as Δ*_k_* (*k* = 1,…,13): The lower and upper boundaries of Δ*_k_* are given as 5(*k* – 1) and 5k, respectively (unit is Å). Table S2 presents the slice range exactly. As discussed in Section 7 of SI and figure S13, the boundary between slices Δ_4_ and Δ_5_ coincides well to the gate of the binding pocket, which was introduced as the boundary between the blue-colored contour region (*ρ_CMb_*(***r**_cube_*) ≥ 0.1) and the magenta-colored contour region (*ρ_CMb_*(***r**_cube_*) ≥ 0.01) in figure 2. By this agreement, four slices Δ_1_,…, Δ_4_ are in the binding pocket, and slices Δ_5_,…, Δ_13_ are outside the pocket.

Figures S14–S17 present conformations picked from some slices and Section 8 of SI explains the features of those picked conformations. These figures visually propose that the contact site on the N-terminal tail to bosentan tends to move with *r_bb_* moving. To check this tendency quantitatively, we introduced a “residue-based bosentan–tail contact ratio” 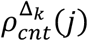, which is the contact ratio of residue *j* of the N-terminal tail to bosentan in a slice Δ*_k_*. Then, we converted 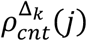 to a potential of mean force (FMF) *F*^Δ*_k_*^(*j*) (equation S14 of Section 9 of SI). Imagine a 2D plane constructed by the residue ordinal number *j* and the slice Δ*_k_*. The lower the *F*^Δ*_k_*^(*j*) at a site [*j*, Δ*_k_*] in the 2D plane, the stabler the contact.

Figure 5 demonstrates *F*^Δ*_k_*^(*j*) in the 2D plane as the free-energy landscape. This figure indicates existence of two free-energy basins, which are denoted by *H*_1_ and *H*_2_ in the figure. We discuss the binding process assuming that bosentan is approaching hETB from a far solvent region to the binding pocket. In Δ_13_ (60 Å ≤ *r_bb_* < 65 Å), the N-terminal tail does not touch bosentan: 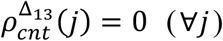. In Δ_12_ (55 Å ≤ *r_bb_* < 60 Å), bosentan first touch Ace79 of the N-terminal tail. Because the system moves generally toward a low free-energy region, possible transition paths are shown by black arrows in figure 5, which lead the conformation to basin *H*_1_. Those motions make *r_bb_* decreased and the bosentan–tail contact firm, resulting in decrement of the free energy.

**Fig. 5.**
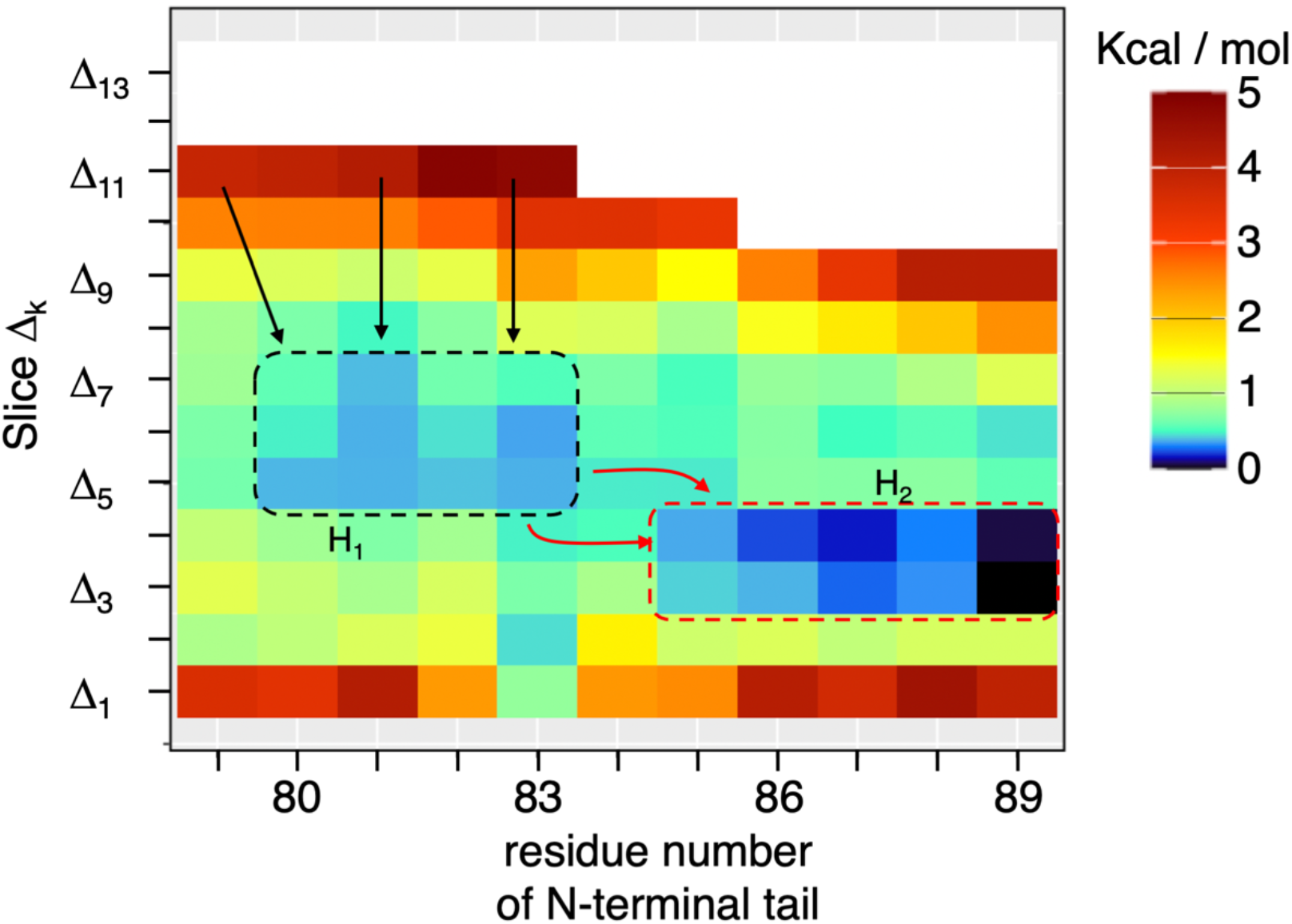
Residue-based bosentan–tail contact ratio, 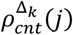, converted to 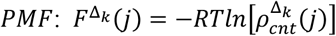 (see Section 9 of SI), where *j* is residue ordinal number (x-axis) of hETB’s N-terminal tail, and Δ*_k_* is the *k*-th *r_bb_* slice (y-axis), where the lower and upper boundaries are 5(*k* – 1) and 5*k*, respectively. Unit is Å. *PMF* value is given by color (see color bar). White is assigned to unsampled regions or regions whose *PMF* are larger than 5 kcal / mol. Sequence of the N-terminal tail is: Ace79, Ser80, Pro81, Pro82, Arg83, Thr84, Ile85, Ser86, Pro87, Pro88, Pro89; Ace is the acetyl group introduced to cap the N-terminal tail tip. Δ*_k_* is range of *r_bb_* slice given in Table S2. Region *H*_1_ ranges from Ser80 to Arg83 in x-axis and from Δ_7_ to Δ_5_ in y-axis. *H*_2_ does from Ile85 to Pro89 and from Δ_4_ to Δ_3_. Arrows are mentioned in text.

With decreasing Δ*_k_* further, the stable region moved from the basin *H*_1_ to *H*_2_, that is, the system moves from the tip region to the root region of the N-terminal tail. This move is natural because *H*_2_ is lower than *H*_1_. Remember that the boundary between Δ_5_ and Δ_4_ corresponds to the gate of binding pocket. Conformational changes from *H*_1_ to *H*_2_ can occur via multiple pathways indicated by the red-colored arrows in figure 5. Because the sites, to which the red arrows are assigned, had a larger free energy than *H*_1_ and *H*_2_ do, the *H*_1_-*H*_2_ basin switch requires overcoming the free-energy barrier.

In Δ_2_ and Δ_1_, which are located at the bottom of the binding pocket, the bosentan–tail contact is disrupted. This does not mean that the complex itself is dissociated but that the complex is stabilized without the bosentan–tail contact, because the native-like complex conformations are stabler than the other state as shown in figures 2 and 3. We also observed a specific contact at Arg83 in Δ_2_ and Δ_1_. However, this specific contact is a weak interaction because the depth of free energy at Arg83 is shallow (figure 5).

### Bosentan–tail non-specific contacts

Figure 5 does not show whether the bosentan–tail contact is specific or non-specific. To investigate the bosentan’s atom specificity, we calculated an “atom–residue contact ratio” 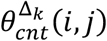 which is the contact ratio of a heavy atom *i* of bosentan contacting to residue *j* of the N-terminal tail in each slice Δ*_k_*. The exact definition of 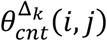 is presented in Section 9 of SI. Figure 6 illustrates 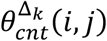 in each slice. The normalization is done for each panel of figure 6 to highlight the atom-specificity (color graduation) in each slice. Thus, the existence of basins *H*_1_ or *H*_2_ is not recognized in figure 6.

**Fig. 6.**
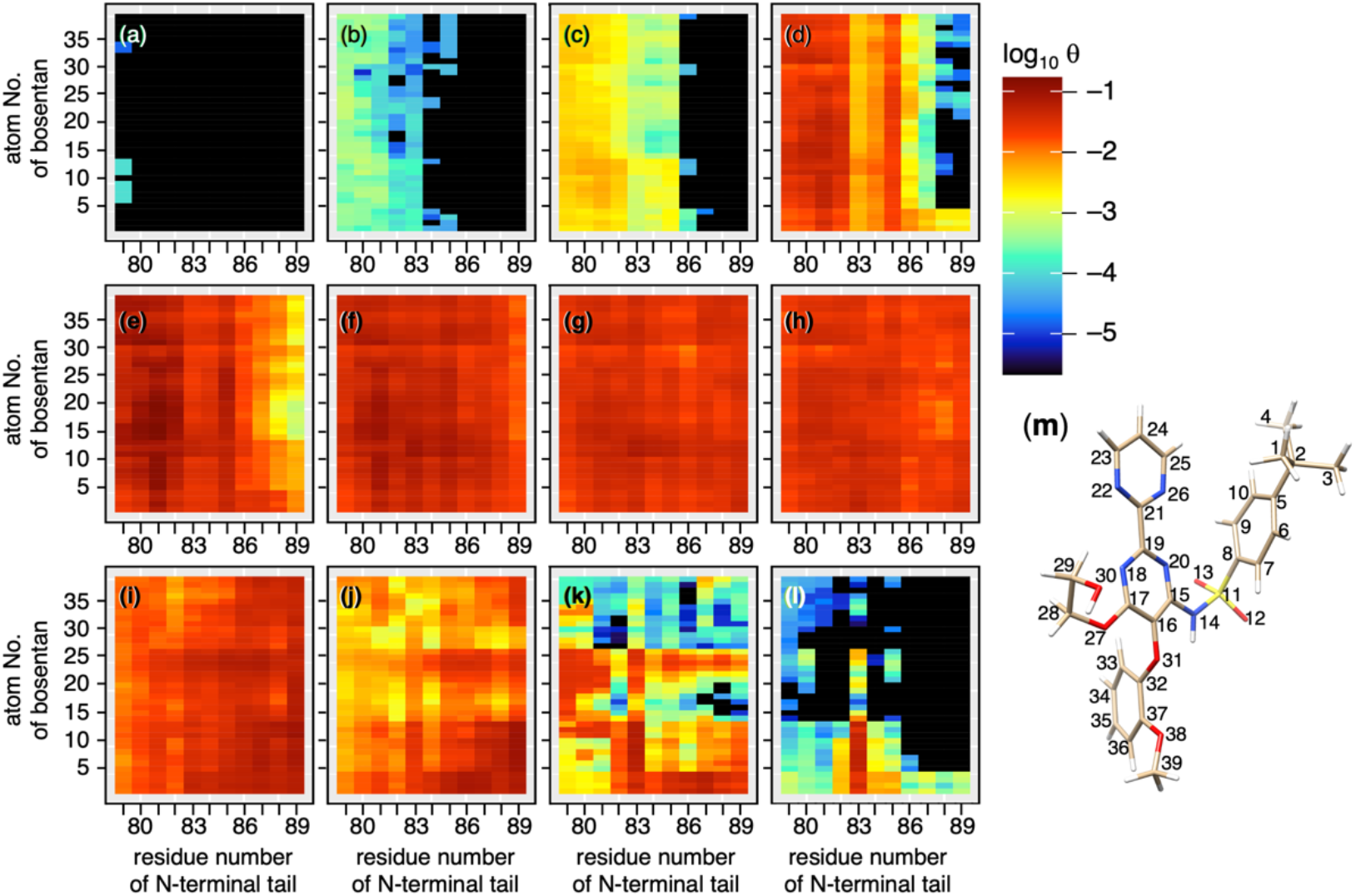
Atom–residue contact ratio 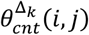, which is contact ratio of heavy atom *i* of bosentan to residue *j* of N-terminal tail in each slice Δ_*k*_. Its exact definition is given in equation S15 of SI. Panels ***a***,…, ***l*** plot 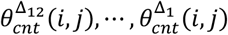, respectively.

Figure 6**a** demonstrates that a few atoms of bosentan contacts rarely to Ace79 of the N-terminal tail in Δ_12_. Figures 6**b**–**d** indicate that all the heavy atoms of bosentan contact non-specifically to the tip of the N-terminal tail in Δ_11_-Δ_9_. Figures 6**e**–**h** show again that bosentan’s contact is non-specific in Δ_8_ – Δ_5_, where all residues of the N-terminal tail contact to bosentan. Up to here (Δ_12_ – Δ_5_), bosentan is outside of the binding pocket. The non-specific contact is still found in Δ_4_ (figure 6**i**) and Δ_3_ (figure 6**j**), although Δ_4_ and Δ_3_ are in the binding pocket.

In Δ_2_ and Δ_1_ (figures 6**k** and **l**, respectively), contrarily, Arg83 of the N-terminal tail contacts specifically to atoms 1–13 of bosentan (see figure 6**m**). This specificity is consistent with figure 5. Note that the atoms 1–10 and 11–13 are hydrophobic and hydrophilic, respectively. Figure S18 demonstrates some conformations that involves the specific bosentan–tail contacts. Generally, the hydrophobic sidechain stem of Arg83 of the tail and the hydrophobic atoms 1–10 of bosentan contact, and the nitrogen atoms of the sidechain tip of Arg83 and oxygen atoms of sulfonamide of bosentan do.

As shown in figures 4 and 5, the bosentan–tail contact tends to be dissociated when bosentan proceeds to the bottom of the binding pocket (Δ_1_ and Δ_2_). Thus, in many snapshots in Δ_1_ and Δ_2_, the N-terminal tail does not contact to bosentan. If the bosentan–tail contact has a role to capture bosentan in solution, we consider that the contact in the binding pocket has no role for molecular binding.

### Fly-casting with ligand sliding

Here we check whether the bosentan–tail contact acts as an attractive interaction. First, we calculated a probability distribution function *P*_Δ_*k*__(*r_b–N_*) regarding the minimum heavy atomic distance, *r_b–N_*, from bosentan to the N-terminal tail for snapshots, which are involved in a slice Δ*_k_*. Then, FMF regarding *r_b–N_* is defined as: *PMF*_Δ_*k*__(*r_b–N_*) = –*RT*ln [*P*_Δ_*k*__(*r_b–N_*)].

Figure 7 presents *PMF* for each slice. Apparently, the bottom of the lowest free-energy basin is assigned to a noncontact state in slices Δ_12_–Δ_10_ (*r_bb_* ≥ 45 Å) (figures 7**a**–**c**). FMF for Δ_13_ is not shown because a bosentan–tail contact was not detected in Δ_13_. In contrast, the lowest free-energy basin is assigned to a contact state (*r_b–N_* ≈ 3.5 Å) in Δ_9_–Δ_2_ (figures 7**d**–**k**), which means that the bosentan–tail contact acts effectively as an attractive interaction in the range of 45 Å > *r_bb_* ≥ 5Å.

**Fig. 7.**
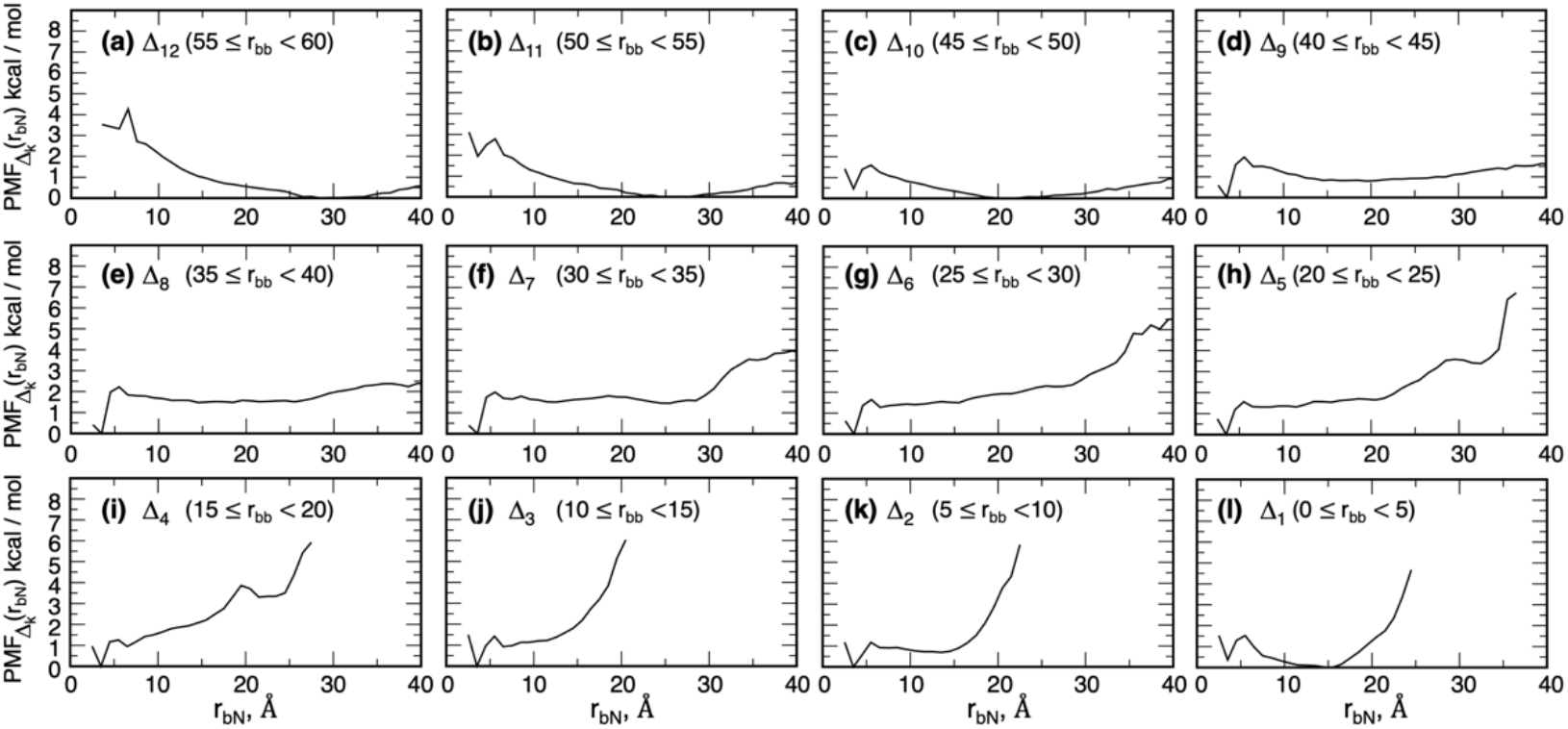
Potential of mean force *PMF*_Δ_*k*__(*r_b–N_*) regarding bosentan–tail distance *r_b–N_* in slice Δ*_k_* (*k* = 1,… 12). *PMF*_Δ_*k*__(*r_b–N_*) is defined in main text. Panels ***a***,…, ***l*** plot *PMF*_Δ_12__(*r_b–N_*),…, *PMF*_Δ_1__(*r_b–N_*), respectively. *PMF*_Δ_13__(*r_b–N_*) is omitted because *PMF*_Δ_12__(*r_b–N_*) is similar with *PMF*_Δ_12__(*r_b–N_*).

Above analysis showed that a bosentan–tail attractive interaction exerted from Δ_9_ (45 Å = *r_bb_*) between the N-terminal tail and bosentan. Together with the residue sliding from the tip side to the root side with decreasing *r_bb_*, we call this binding mechanism “fly-casting with ligand sliding”.

### Bosentan–membrane Contact

Bosentan occasionally contacts to membrane (figures S14**e**, S15**d** and S15**e**). To investigate the bosentan–membrane contact, we introduced a quantity, a “bosentan– membrane contact ratio” 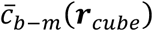, which is calculated with a similar procedure with 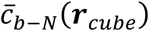: See Subsection 3.3 of SI for details of the calculation procedure. The bosentan–membrane contact was formed commonly on the surface of the membrane (figure S19). Some experiments^44,45^ suggest a possibility that the ligand sinks and diffuses in the lipid bilayer to reach the binding pocket of GPCR rather than approaching the pocket from the solvent side. In the present simulation, however, we could not find such a ligand diffusive motion in membrane, whereas we observed bosentan sank in membrane (Figure S14**e**). Thus, we conclude that bosentan mainly approach the pocket from solvent. Because membrane prevents bosentan from moving to cytoplasmic side, the membrane has a role to restrict the phase volume for bosentan to move. This volume restriction may facilitate binding of bosentan to the binding pocket.^46^

### Orientational selection when bosentan entering the binding pocket

We investigated the molecular orientation of bosentan around hETB. First, we defined the bosentan’s molecular orientation by two vectors (orientation vectors) ***v***_↓_ and ***v***_←_, which are fixed in the core region of bosentan (figure S20**a**), which are approximately perpendicular to each other (see figure 3**a** for exact definition of the core region). Then, we normalized the orientation vectors as |***v**_α_*| = 1 (*α* =← or ↓). Because the bosentan’s core region is structurally stiff, the two orientation vectors are convenient quantities to define the bosentan’s molecular orientation. We note that in general two nonparallel vectors can specify the orientation of a plane (the central ring of the core region) in the 3D real space. Figure S20**b** exemplifies the vectors in the native complex and in a snapshot. Next, we assigned the orientation vectors to a cube ***r**_cube_*, in which the bosentan’s centroid was detected. Then, Ω in equation S7 of Subsection 3.4 of SI is replaced by ***v**_α_*, and the resultant quantity 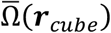 is referred to as 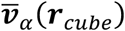 in the equation. We named 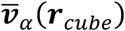 as an “averaged orientation vector”. Because ***v**_α_* in a snapshot was normalized, the maximum of 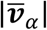 is unity when all the vectors detected in the cube *r_cube_* are perfectly ordered, and the minimum is 0.0 when the vectors are completely randomized.

Figures 8**a** and **b** illustrate the spatial patterns of 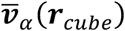, in which orientation vectors with relatively large norms 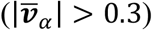 are shown. Importantly, the large-norm vectors were found mainly in the binding pocket. Therefore, the bosentan’s orientation is relatively ordered in the pocket whereas it is randomized outside the pocket. This result suggests that a configurational entropy of bosentan decreases quickly when bosentan passes the gate of the binding pocket. Remember that the free-energy basin of *F*^Δ_*k*_^(*j*) also switched quickly from *H*_1_ to *H*_2_ at the gate (figure 5). Thermodynamically the decrease of entropy results in increase of free energy. This entropic decrease is compensated by another thermodynamic factor, otherwise, the density *ρ_CMb_*(***r**_cube_*) should decrease in the binding pocket. We show in the next section that intermolecular native contacts (attractive electrostatic interactions) are formed in the binding pocket. This can enthalpically compensate the free-energetic disadvantage by the entropy decrement brought by the loss of entropy.

**Fig. 8.**
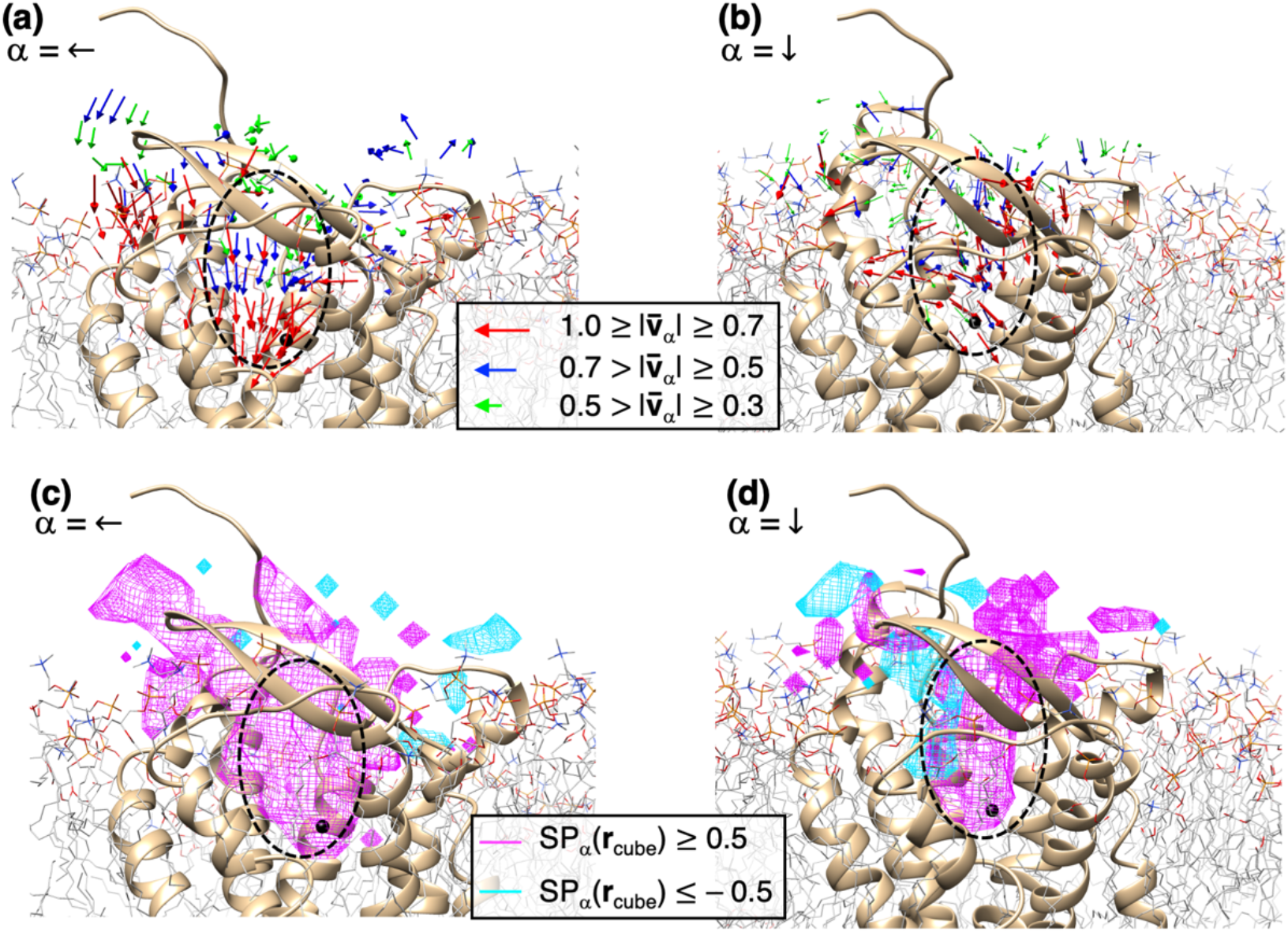
Spatial patterns of averaged orientation vectors (**a**) 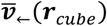 and (**b**) 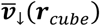. Vector is colored differently by its norm 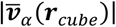 (*α*= ← or ↓) (see inset). Vectors with 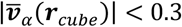 are omitted. (**c**) Spatial patterns of scalar product 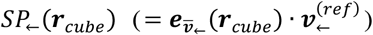 and (**d**) those of 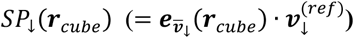, where 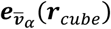 and 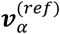 are defined in main text. Regions of *SP_α_*(***r**_cube_*) ≥ 0.5 and *SP_α_*(*r_cube_*) ≤ −0.5 are colored differently by their values (see inset). For all panels, structure of hETB is from native complex, and small black spheres are bosentan’s centroid in the native complex. Binding pocket is shown by broken-line circle in all panels.

Figures 8**a** and **b**, however, do not necessarily mean that the bosentan’s orientations are like those in the native-complex position. Then, we check the orientation similarity as follows: First, we calculated ***v***_←_ and ***v***_↓_ for the native complex, which are referred to as 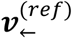 or 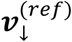, respectively. Next, we calculated a scalar product as: 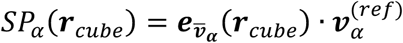, where 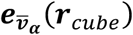 is a unit vector parallel to 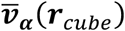. If 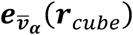 is exactly parallel to 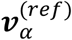, then *SP_α_*(***r**_cube_*) = 1, and if 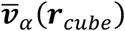 is exactly anti-parallel to 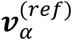, then *SP_α_*(***r**_cube_*) = −1. If 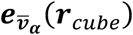 and 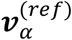 are perpendicular to each other, then *SP_α_*(***r**_cube_*) = 0. The spatial pattern of *SP_α_*(***r**_cube_*) is a measure to assess the similarity between 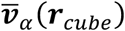 and 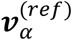.

Figures 8**c** demonstrates that the major volume of the binding pocket was occupied by *SP*_←_(***r**_cube_*) ≥ 0.5, which means that the orientations of 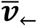 and 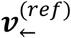 were similar. For 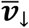, a large volume of the pocket was also characterized by *SP*_↓_(***r**_cube_*) ≥ 0.5 whereas a minor part (cyan-colored contours) of the pocket was characterized by *SP*_↓_(***r**_cube_*) ≤ −0.5 (figure 8**d**). This minor part does not cover the bosentan’s native position at all. Therefore, we conclude that bosentan enters the pocket with limited orientations advantageous for reaching the native-complex position because the pocket is too narrow for bulky bosentan to flip. We call this possible mechanism “orientational-selection.”

### Bosentan–hETB contacts and ring-C packing at the bottom of binding pocket

Up to here, we analyzed the approaching process of bosentan to the binding pocket of hETB. In this section, we investigate the interactions of bosentan to amino-acid residues of hETB in the binding pocket.

First, we observe the intermolecular interactions in the three X-ray structures, the bosentan–hETB^10^, ET1–hETB^6^, and the K8794–hETB^10^ complexes. We note that the same binding scheme is seen in the three complex structures (figure S21). By viewing the three complex closely, we found four inter-molecular native atomic contacts, which exist commonly in the three complex structures as attractive interactions (See Section 10 of SI and figure S22): Contact (i) between oxygen atoms in sulphonyl group (SOO) of the bosentan’s sulfonamide and the nitrogen (*N_ζ_*) of Lys 182, (ii) between the oxygens in bosentan’s SOO and the nitrogen of Lys 273, (iii) between the oxygens in bosentan’s SOO and nitrogen atoms of Arg 343, and (iv) between the nitrogen atom (*N*14) of the bosentan’s sulfonamide and the nitrogen (*N_ζ_*) of Lys 182.

To analyze the inter-molecular distances of snapshots, we introduce a notation *r_α;β_*, which is a distance between atoms *α* and *β* belonging to the ligand (bosentan, K8794, or ET1) and hETB, respectively. Then, the four distances regarding the four native contacts are denoted as *r*_N14;Lys182N_ζ__, *r*_SOO;Lys182N_ζ__, *r*_SOO;Lys273N_ζ__, and *r*_SOO;Arg343N_η__. When multiple distances were possible in *r_α;β_*, the minimum of the distances was selected for *r_α;β_*. For instance, four distances are possible for *r*_SOO;Arg343N_η__ because two oxygen atoms and two nitrogen atoms are involved in the sulphonyl group and Arg 343, respectively. Table S3 lists the distances *r_α;β_* in the X-ray structures, in which three distances are around 3 Å except for *r*_SOO;Lys182N_ζ__ (= 4.56 Å) from the bosentan-hETB complex. We presume that *r*_SOO;Lys182N_ζ__, *r*_SOO;Lys273N_ζ__, and *r*_SOO;Arg343N_η__ are related to salt-bridges, and *r*_N14;Lys182N_ζ__ may be done to either a saltbridge or a hydrogen bond. This point is discussed later.

We judged that the heavy atoms *α* and *β* are contacting if *r_α;β_* < 5.0 Å is satisfied in a snapshot. The value of 5.0 Å is about 0.5 Å larger than the largest (4.56 Å) of the experimental values (Table S3). Considering thermal fluctuations of the complex structure and an error in experimental measurement (3.6 Å resolution for the bosentan– hETB complex), we added the tolerance of 0.5 Å to the inequality. Besides, a heavy atom has a radius of 2.0 Å approximately, and the remaining distance between *α* and *β* is 1.0 Å (=5.0 Å – 2 × 2.0 Å), which does not allow a water molecule to penetrate because the diameter of a water molecule is about 3.0 Å. Therefore, the threshold of 5.0 Å is rational to check the atomic contacts.

Based on the criteria above, we calculated the number of contacts *N_cnt_* (0 ≤ *N_cnt_* ≤ 4) regarding the four distances for all snapshots. Then, picking snapshots falling in an *r_bb_*-range of *R*(*r_bb_*) = [*r_bb_* – Δ*r_bb_*; *r_bb_* + Δ*r_bb_*], we calculated average of *N_cnt_* for the picked snapshots, and denoted it as 〈*N_cnt_*(*r_bb_*)〉. Section 11 of SI presents the exact method to calculate 〈*N_cnt_*(*r_bb_*)〉 and its standard deviation 〈*SD_cnt_*(*r_bb_*)〉. Ideally, Δ*r_bb_* should be sufficiently small (Δ*r_bb_* → 0) to calculate 〈*N_cnt_*(*r_bb_*)〉 exactly at *r_bb_*. However, the number of snapshots in *R*(*r_bb_*) decreases with decreasing Δ*r_bb_* because the simulation length is finite. To maintain the statistical significance in 〈*N_cnt_*(*r_bb_*)〉, we set Δ*r_bb_* = 0.25 Å for the present simulation. 〈*N_cnt_*(*r_bb_*)〉 is 4 if all snapshots in *R*(*r_bb_*) always have four contacts and zero if all snapshots always have no contact.

Figure 9 depicts the *r_bb_* dependence of 〈*N_cnt_*(*r_bb_*)〉, where *r_bb_* starts from 0.25 Å and 〈*N_cnt_*(0.25 Å)〉 is calculated for range of *R*(0.25 Å) = [0.0 Å; 0.5 Å]. *R*(*r_bb_* < 0.25 Å) is impossible to be calculated. We set the maximum of *r_bb_* at 15.25 Å, where the used range is *R*(15.25 Å) = [15.0 Å; 15.5 Å]. Note that *R*(15.25 Å) is involved in the slice Δ_4_, just below the gate of the hETB’s binding pocket (figure S13). Therefore, figure 9 shows that almost no atom contacts are formed at the gate: 〈*N_cnt_*(15.25 Å)〉 = 0.005. 〈*N_cnt_*(*r_bb_*)〉 increased with decreasing *r_bb_* (with bosentan approaching the native-complex position): The number of contacts exceeded the value of 1 at *r_bb_* = 3.25 Å: 〈*N_cnt_*(3.25 Å)〉 = 1.03 ± 0.85. Subsequently, the growth rate of 〈*N_cnt_*(*r_bb_*)〉 rose, and 〈*N_cnt_*(*r_bb_*)〉 reached the maximum at *r_bb_* = 0.25 Å: 〈*N_cnt_*(0.25 Å)〉 = 2.74 ± 0.79. We further investigated formation of each contact (see Section 12 of SI). Figure S23 indicates that each native contact is formed with decreasing *r_bb_*.

**Fig. 9.**
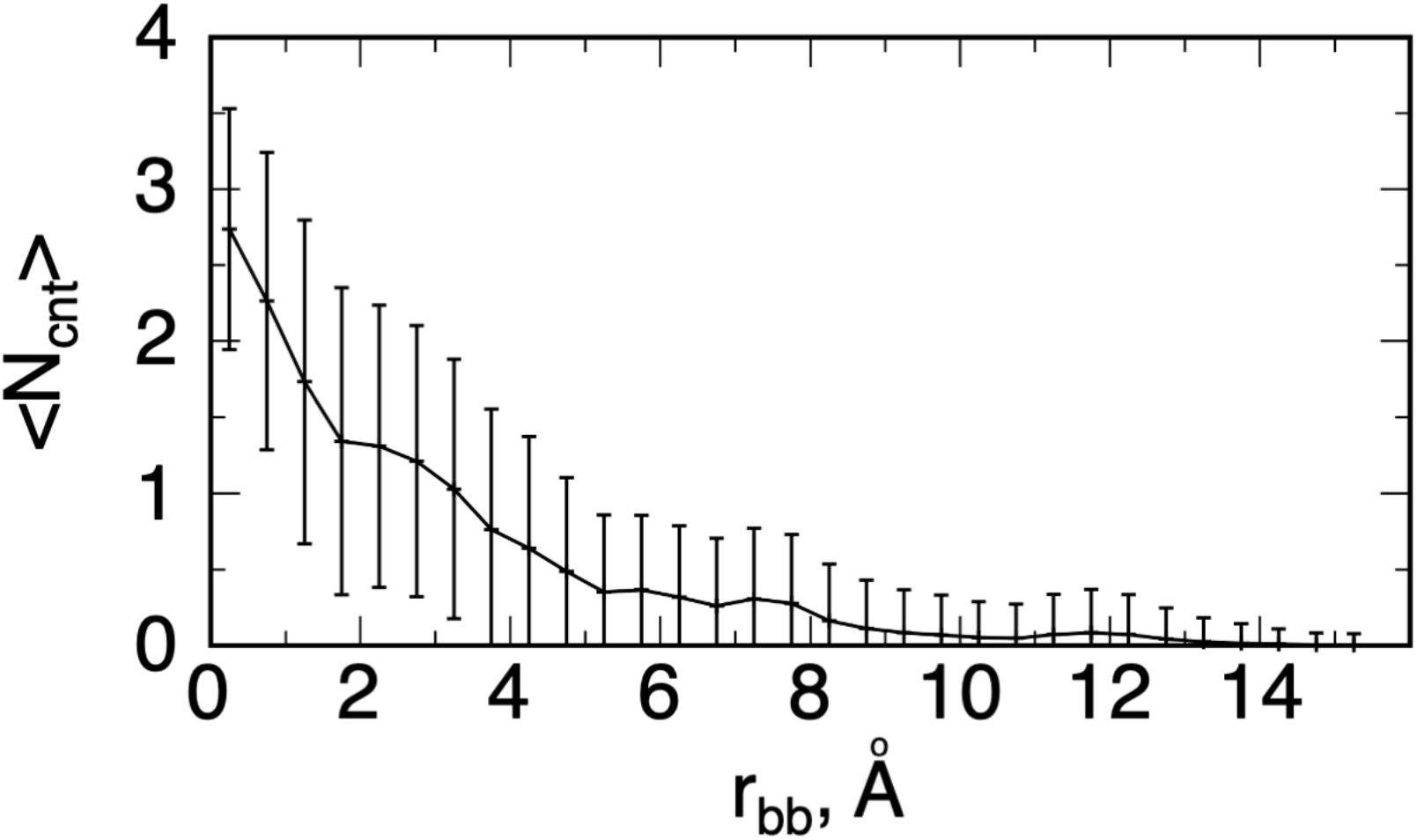
Dependency of 〈*N_cnt_*(*r_bb_*)〉 on *r_bb_* and its standard deviation (error bars). Lower limit of some error bars is smaller than 0 on the y-axis because the standard deviation was calculated using usual form: [〈*N_cnt_*(*r_bb_*)^2^〉 – 〈*N_cnt_*(*r_bb_*)〉^2^]^2^.

Because the four intermolecular native contacts act as attractive interactions (Section 10 of SI), figure 9 indicates that the complex structure is stabilized enthalpically by the native contacts when bosentan goes to the bottom of the binding pocket. We presume that the instability induced by the entropy decrement when bosentan passing the gate of the binding pocket is compensated by the formation of the native contacts.

We checked the reproducibility of the packing of ring C to the bottom of the binding pocket (see figure S21**c** for ring C of bosentan). For this purpose, we calculated the density map, *ρ_CMC_*(***r**_cube_*), of the centroid of the ring C using equation S3 of Subsection 3.1 of SI with replacing the centroid of portion *X* by the centroid of ring *C*. Figure S24 demonstrates the high-density spot (*ρ_CMC_*(***r**_cube_*) ≥ 0.8) for ring C. Importantly, the high-density spot is localized in a region that overlaps with the ring-C position in the native complex. On the other hand, the contours of *ρ_CMC_*(***r**_cube_*) ≥ 0.4 shows a deviation from the native-complex position. We discuss this discrepancy between the computation and the native complex later.

### Bosentan’s conformations in the highest-density spot

As shown in figure 2, the bosentan’s centroid in the native complex structure was located on the fringe of the computed highest-density spot (red-colored contours in Figure 2). In this section, we view the bosentan’s conformations in the highest-density spot.

Remember that the highest-peak position of 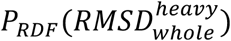 or 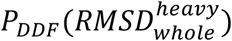 of figure 3 was larger than that of 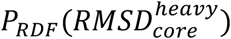 or 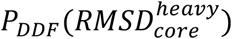. This suggests that the sidechains of bosentan were somewhat disordered. To examine this disorder in more detail, we collected snapshots from 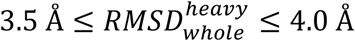, which involves the highest peak of 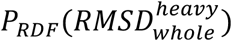 (Figure 3**b**).

Figure 10**a** displays four conformations randomly pocked from the snapshots collected above. This figure demonstrates that the conformations from the highest density spot have a similarity with the native-complex structure (the X-ray structure). Thus, we referred to the complex conformations in highest density spot as “native-like complex”. On the other hand, Figure 10**a** indicates that the longest sidechain of bosentan, ring A (Figure S21**c**), is deviated from the native-complex position with a conformational diversity, whereas the core region was relatively converged to a position close to the native-complex position. This is why the highest-peak position of 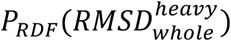 or 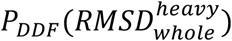 was larger than that of 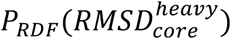 or 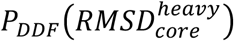 in figure 3. We discuss the imperfectness of ring A in the next section.

**Fig. 10.**
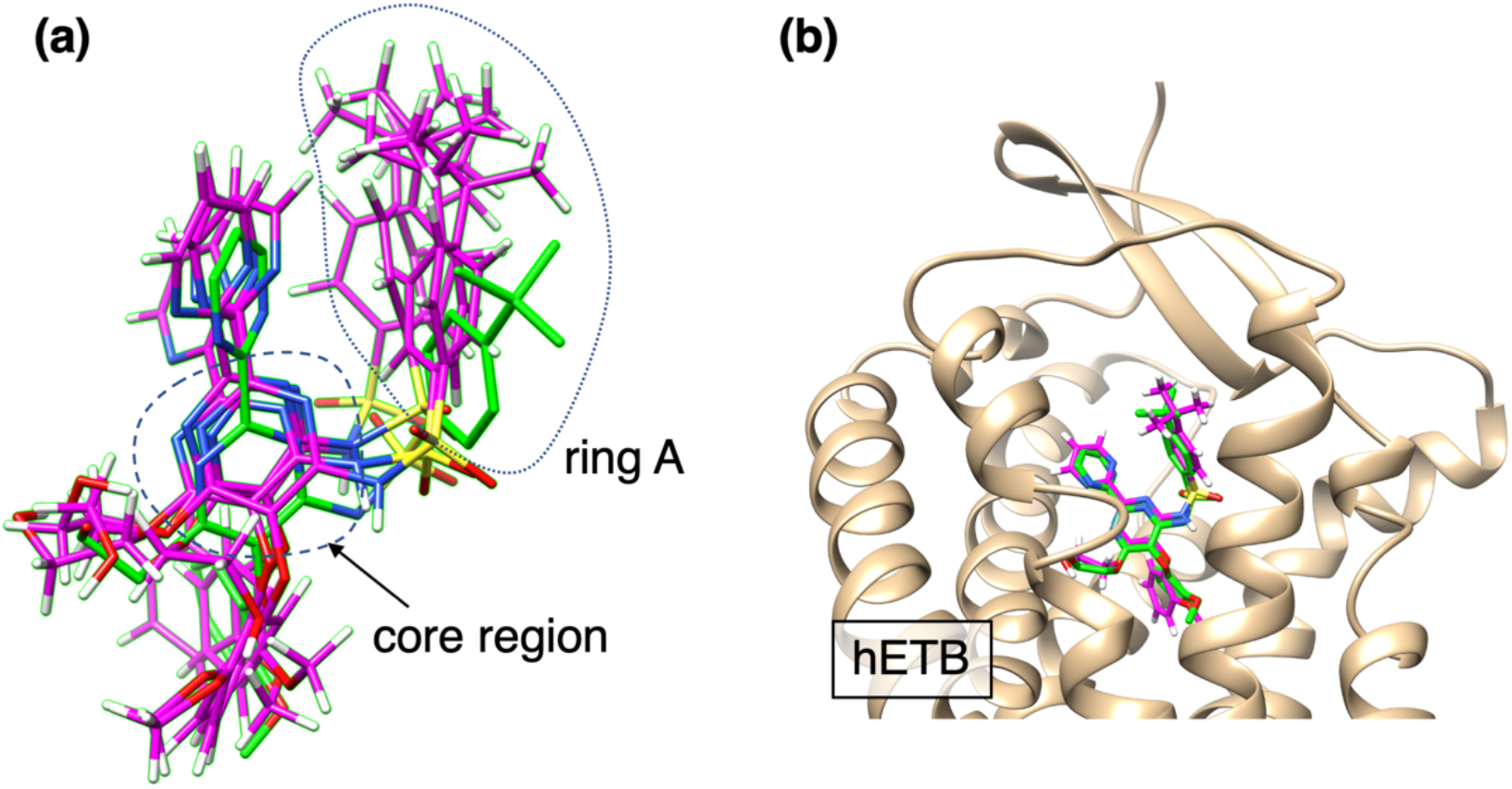
Snapshots sampled in the highest density spot (*ρ_CMb_*(***r**_cube_*) = 0.5 in figure 3). In both panels, bosentan’s conformations from simulation are shown in magenta, and one in the native complex is shown in green. (**a**) Four conformations picked randomly from range of 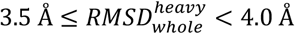. Bosentan’s core region and ring A are shown in figure. (**b**) A conformation with 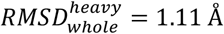. hETB in the native complex is also shown. Note that superimposition was not applied to bosentan but done to transmembrane helices of hETB (see figure S7**a**).

Figure 10**b** illustrates a snapshot with a small 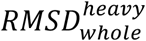 (1.11 A). Such small 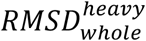 conformations were minority in the ensemble of snapshots because there was no peak at 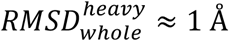 in figure 3. This small 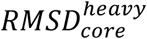 was resulted because the ring A of this conformation overlapped well on that of the native-complex structure.

**Fig. 10.** Snapshots sampled in the highest density spot (*ρ*_CMb_(***r**_cube_*) = 0.5 in figure 3). In both panels, bosentan’s conformations from simulation are shown in magenta, and one in the native complex is shown in green. (**a**) Four conformations picked randomly from range of 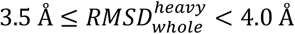. Bosentan’s core region and ring A are shown in figure. (**b**) A conformation with 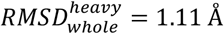. hETB in the native complex is also shown. Note that superimposition was not applied to bosentan but done to transmembrane helices of hETB (see figure S7**a**).

### Comparison of bosentan at the highest-density spot with that in the crystal complex structure

We note several discrepancies between bosentan in the crystal complex structure and that at the computed highest-density spot. GA-guided mD-VcMD is powerful enough to explore the ligand–receptor conformational space starting from completely unbound conformations (figure S5). However, we remark that reproduction of the native-like complex was not perfect. As discussed, the bosentan’s longest sidechain (ring A; figure S21**c**) exhibited a structural diversity in the native-like complex conformations although the core region agreed well with the native-complex position.

We presume two possibilities for the imperfectness: Accuracy of classical-mechanical force-fields and/or treatment of the bosentan’s sulfonamide. Despite lots of attempts for improving the force fields, no perfect ones have not yet obtained as well known. In future, novel force-field parameters may solve the detailed conformational inconsistency with experiment.

Regarding the second possibility, we used the bosentan’s sulfonamide in a normal (neutral) form in that a hydrogen atom is attached to the nitrogen atom (N14) of the sulfonamide. In the neutral form, the electronic state of N14 is the sp^2^ hybridization with planner three-chemical bonds (S-NH-C angle ~130 degrees). In the deprotonated form, the electronic state is also the sp^2^ hybridization with planner two-chemical bonds and one electron-lone pair (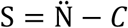 angle ~131 degrees). Therefore, we cannot distinguish the electronic state of sulfonamide by viewing the X-ray structure of bosentan. We may treat the N14 atom deprotonated because the *pKa* value of bosentan is 5.1 in solution^47^. However, the form of sulfonamide in the binding pocket of hETB is unknown. There is a report^48^ that *pKa* of sulfonamide in various compounds, which are immersed in mixture of organic solvent and water, increases largely with increasing the fraction of the organic solvent. Also, as reported^49^, the electronic state of aminoacids in the protein interior may be different from that in solution. Finally, we decided to use the normal form of bosentan with protonated N14. We note that the distribution function *ρ_bs_*(*r*_*N*14;*Lys*182*N*_ζ__;*r_bb_* = 0.25 Å) exhibited a remarkable peak at *r*_*N*14;*Lys*182*N*_ζ__ = 3 Å (figure S23**f**), reproducing the experimental observation (Table S3). This result supports that the currently used form of sulfonamide is plausible. To analyze further the structure diversity of the ring A, huge computations are required with varying the force-fields and the charge state of sulfonamide in future.

## Conclusions

Prediction of the ligand–receptor complex structure is important for structure-based drug design, and the recent fast MD simulations have enabled the ligand–receptor docking MD simulations. This strategy is reasonable to develop a new drug (especially antagonist) because the molecular system finally reaches the most stable structure regardless of the binding pathways. However, the binding process is also important to understand the complex. The current simulation study showed that an attractive interaction acts effectively between bosentan and the N-terminal tail of hETB via the fly-casting mechanism^17,18,19^ even when bosentan is located distant from the binding pocket of hETB. Some experiments have reported that the N-terminal tails of GPCRs have important physiological functions^50^ whereas the N-terminal tails of GPCRs have been studied insufficiently in contrast to the binding pocket. Because the tail is conformationally disordered, a simulation like this study is helpful for understanding the function or role of the N-terminal tail.

In general, GPCRs have a long and disordered N-terminal tail (40–200 aminoacid residues long)^21^. An MD study showed that an amino-acid substitution to the N-terminal tail of a GPCR (human *β*_2_-adrenergic receptor) varies the dynamics of the tail, and that the accessibility of the ligand to the binding pocket also varies by the substitution^51^. This study as well as our present work suggest that the ligand–receptor interaction should be studied even when the ligand is outside the binding pocket of GPCR.

The bosentan–hETB complex formation obtained from the present study is summarized as follows: First the tip-side of the N-terminal tail of hETB captures bosentan when bosentan is fluctuating in solution via nonspecific intermolecular contacts. The bosentan–capturing conformation is identified as the free-energy basin *H*_1_ in the free-energy landscape (figure 5). Then, the ligand–sliding occurs occasionally from the tip-side to the root-side of the N-terminal tail when bosentan passing the gate of the binding pocket, which corresponds to basin switching from *H*_1_ to the deeper basin *H*_2_ with overcoming the inter-basin free-energy barrier. We named this mechanism “fly-casting with ligand–sliding”. Importantly, the molecular orientational variety of bosentan decreases quickly when bosentan passing the gate (figure 8). This decrement of the orientational variety has a role of screening out bosentan’s conformations whose molecular orientations are improper to reach the binding site (orientational selection). We note that this orientational selection is a variant of conformational selection^20^. Because the decrement of the bosentan’s orientational variety results in loss of a configurational entropy of bosentan, another thermodynamic quantity should compensate the free-energetic disadvantage by the decrement of the orientational variety. Otherwise, the density of the bosentan’s centroid, *ρ_CMb_*(***r**_cube_*), should decrease in the binding pocket. Our results show that the intermolecular native contacts (electronically attractive interactions) between bosentan and hETB are formed in the binding pocket (figure 9). It is likely that this native contact formation is the compensation factor, and eventually, the native-complex conformation is completed.

We did not use the full-length N-terminal tail for the simulation, whereas we increased the tail length than the previous study^12^: Ten residues longer than the N-terminal residues in the X-ray crystal structure for the present study, and five residues longer than those in our previous study. The present study suggests a possibility that a longer N-terminal tail adds a new property to the binding mechanism. In fact, it has been reported that long N-terminal tails exert various physiological functions of GPCRs such as ligand recognition, receptor activation, and signaling^22,23^. Thus, a simulation with a longer N-terminal tail is the next challenge for the ligand–GPCR simulation.

The shape of the resultant free-energy landscape (potential-of-mean-force surface) was similar between the present and previous studies (figure S10 and Figure 4A of ref. 12). However, the scale of the free energy was considerably different between the two studies: The free-energy slope from the previous study was ten times steeper than that from the present study. Previously, we inferred that the steep slope from the previous study is probably caused by a drawback in the update method of 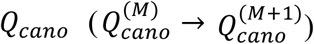 and that the steep slope would be weakened if two pieces of 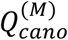, which are assigned to neighboring zones in the RC space, are connected adequately^12^. GA-mD-VcMD is such a method to connect 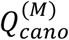 smoothly^13^. In fact, the free-energy slope decreased in the present study.

## Acknowledgements

This work was supported by JSPS KAKENHI Grant No. 21K06052 (J.H.), 20H03229 (N.K.), and JP18H05534 (H.K.), and was performed in part under the Cooperative Research Program of the Institute for Protein Research, Osaka University, CR-21-05. J.H., K.K. and K.N. acknowledge support by the HPCI System Research Project (Project IDs: hp200025, hp200063, hp200090, hp210002, hp210005, and hp210008). J.H., S.I., N.H., I.F., N.K., and Y.F. are supported by the Development of innovative drug discovery technologies for middle-sized molecules project from Japan Agency for Medical Research and Development (AMED). J.H., N.K., I.F., and Y.F. are supported by Project Focused on Developing Key Technology for Discovering and Manufacturing Drugs for Next-Generation Treatment and Diagnosis (2018-2021 and 2021-) from AMED. J.H., K.K., S.S. and H.K. were partially supported by Platform Project for Supporting Drug Discovery and Life Science Research (Basis for Supporting Innovative Drug Discovery and Life Science Research (BINDS)) from AMED under Grant Number JP21am0101106. Authors thank Dr. Isao Yasumatsu from Daiichi Sankyo RD Novare Co., Ltd (Japan) for helpful discussion. GA-mD-VcMD was performed on the TSUBAME3.0 supercomputers at the Tokyo Institute of Technology.

## Author contributions

All authors contributed equally.

