## Supplementary Information (SI) for "Fly-casting with ligand–sliding and orientational selection to support the complex formation of a GPCR and a middle-sized flexible molecule"

#### Section 1. Definition of a reaction coordinate (RC)

Consider two atom groups $G_h^A$ and $G_h^B$ $(h = \alpha, \beta, \gamma)$ in a molecular system. A single reaction coordinate (RC) $\lambda^{(h)}$ is defined by the distance between the centroids of $G_h^A$ and $G_h^B$ (figure S1). Superscripts $A$ and $B$ indicate simply that the two atom groups are pairing to define the single RC, and then, one can exchange the superscripts as: $G_h^A \rightarrow G_h^B$ and $G_h^B \rightarrow G_h^A$ without changing the value of $\lambda^{(h)}$. The actual set of the atom groups for the present system is explained later.

Although we used centroids of atom groups to define RCs in this paper, RCs can be defined arbitrarily by the system's coordinates in theory. To define RCs, for instance, one can use angles that can be described by system's coordinates.

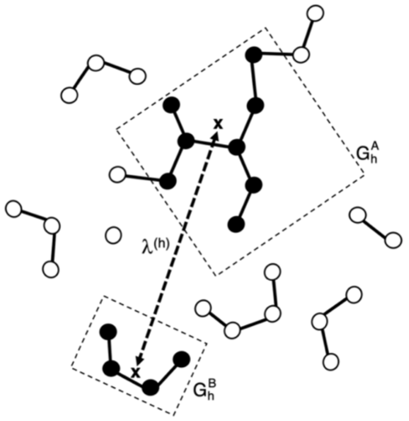


**Figure S1:** Two atom groups, $G_h^A$ and $G_h^B$, are indicated by two rectangles where constituent atoms of the groups shown by black filled circles. Centroid of each atom group is presented by a cross. The distance indicated by broken-line with arrows between the centroids is $\lambda^{(h)}$.

#### Figure S2

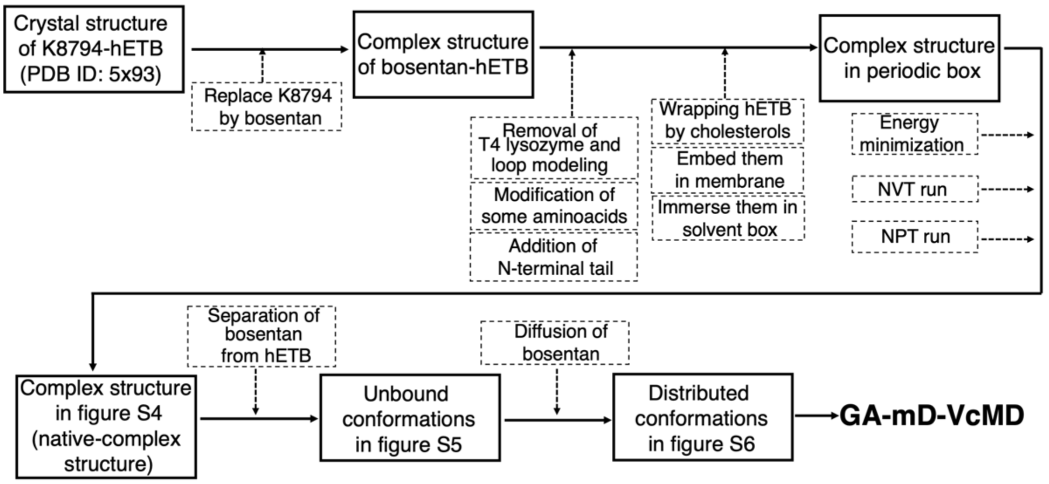


**Figure S2:** Generation of molecular system. A solid-line rectangle represents system at each stage, and solid-line arrow represents conversion of the system from a stage to the next stage. "Complex structure in figure S4", which is close to the X-ray structure of bosentan–hETB complex (PDB ID: 5xpr), may be called the "native-complex structure" in this paper if necessarily. Broken-line rectangles and broken-line arrows indicate actions applied to the system to proceed the stage. "Diffused conformations in figure S6" are initial conformations for GA-mD-VdMD simulations.

#### Section 2. Generation of the molecular system

To generate the bosentan–hETB molecular system, we can use either of two crystallographic complex structures:[1] Bosentan-hETB (PDB ID: 5xpr) and K8794-hETB (PDB ID: 5x93). The chemical compound K8794 is an analog of bosentan. Figure S3**a** illustrates the two complex structures, which are similar to each other. Both compounds bind to the pocket of hETB, and importantly, their positions in the complexes are almost identical (figure S3**b**). K8794 has a long sidechain indicated by a magenta-colored broken-line circle in figure S3**b**, which does not exist in bosentan.

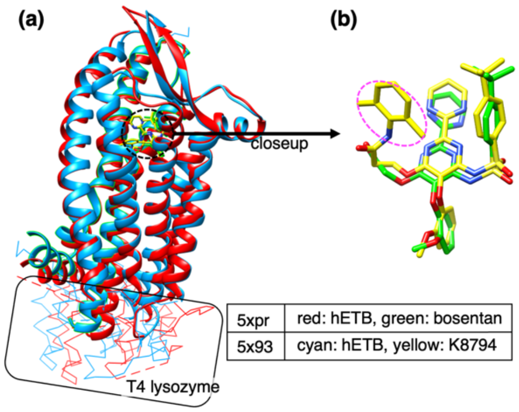


**Figure S3:** (**a**) Superposition of two pdb structures: bosentan-hETB (5xpr) in red and K8794-hETB (5x93) in cyan. Superimposed regions (residues 90–129, 135–206, 217–303, and 311–401) are shown in ribbon models, and $C_\alpha$ atoms in these regions are superimposed. Both structures involve T4 lysozyme (wire model regions surrounded by black solid line), which were artificially introduced to improve crystallogenesis. We removed T4 lysozyme in our simulation because T4 lysozyme does not exist in the native hETB. Black broken-line circle indicates positions of compounds bosentan (green) and K8794 (yellow). (**b**) Closeup of bosentan and K8794 after removing hETB from panel (**a**). Further superposition was not applied to the compounds to draw panel (**b**). Portion indicated by red-colored broken-line circle in magenta exists only in K8794.

We select the K8794–hETB (5x93) complex to generate the computed system because 5x93 has a better resolution than bosentan-hETB (5xpr) complex: 2.2 Å and 3.6 Å for 5x93 and 5xpr, respectively. Then, we replaced K8794 of 5x93 by bosentan. The wild-type hETB consists of 442 amino-acid residues, which are numbered from 1 to 442 in UniPlot (https://www.uniprot.org/uniprot/P24530.fasta). In the crystallographic structure, the intracellular loop 3 (residues 304–310) of hETB was replaced by T4 lysozyme. Then, to generate the genuine hETB molecule, we removed T4 lysozyme and modeled the loop 3. In the crystallography, seven amino-acids of hETB were conducted to increase the thermostability of hETB (R124T, D154A, K270A, S342A, and I381A) or to prevent heterogenous palmitoylation in hETB (C396A and C400A). Then, we back-mutated those amino-acids to the wild-type ones. Besides, some missing atoms in the crystallographic structure were modeled.

Although hETB is originally comprised of residues 1–442, the N-terminal residues 1–84 and the C-terminal residues 405–442 are missing in the X-ray structure of

5x93. In our previous simulation of the ET1–hETB system[2], we treated residues 85–403 explicitly and found that the N-terminal tail (residues 85–89) of hETB captures ET1 prior to binding of ET1 to the pocket of hETB (the fly-casting mechanism[3,4,5]). In the present study, we extended the N-terminal tail up to the 80-th residue to study the fly–casting mechanism further. Therefore, the N-terminal tail consists of residues 80–89 in the current study. We did not extend the C-terminal tail of hETB because the C-terminal tail is in the cytoplasmic side (no direct interaction to bosentan). Finally, hETB consists of residues 80–404 in the simulation.

According to a method explained in our previous paper (Section 8 of SI of ref. 2), we embedded hETB in a membrane (the popc-lipid bilayer) and put four cholesterol molecules in the hETB–membrane interface because these cholesterols play an important role to stabilize the GPCR (i.e., hETB) tertiary structure.[6] Up to here, the system consists of bosentan, hETB, and membrane (cholesterol and POPC lipid molecules).

The system generated above was immersed in a periodic-boundary box filled by water molecules. The sides of the box were parallel to the x-, y-, or z-coordinate axis (the x-y plain was parallel to the membrane surface), where the box size was 71.33 Å (x-axis), 71.33 Å (y-axis), and 132.17 Å (z-axis). Then, we introduced 28 sodium and 43 chlorine ions with replacing water molecules randomly by the ions to set the net charge of the entire system to zero and to set the buffer ionic concentration to a physiological one. The number of the final constituent atoms of the system is 69,062 (numbers of protein and bosentan atoms are 5289 and 68, respectively; number of cholesterols, POPC, and water molecules are 4 (296 atoms), 127 (17018 atoms), and 15440 (46320 atoms), respectively; number of sodium and chlorine ions are 28 and 43, respectively). This conformation is denoted as "Complex structure in periodic box" in figure S2.

After energy minimization of the above system, a short NVT (constant volume and temperature) simulation was performed. Then, an NPT (constant pressure of 1 atm and temperature of 300 K) simulation was performed to equilibrate the box size. The resultant box size was $69.37\text{Å} \times 69.37\text{Å} \times 140.29\text{Å}$. Figure S4 illustrates the resultant conformation, which shows that bosentan is close to the natively bound position because the generation of the molecular system started from the X-ray structure of 5x93. We denote this conformation "Complex structure of figure S4 (native-complex structure)" in figure S2. We emphasize that this complex is not the initial conformation of GA-mD-VcMD: As explained in the main text, two more stags are processed to obtain the initial conformation, where bosentan was completely separated from the binding pocket of hETB.

Last in this section, we note that the conformation of the N-terminal tail in the native complex (figure S4) should be regarded as an instantaneous conformation because the tail is an intrinsically disordered segment, whereas the other part of the hETB is determined in the crystal structure[1]. We report in this paper that the N-terminal tail fluctuates largely during the simulation. This result agrees with the fact that the N-terminal tail is missing in the crystal structure.

#### Figure S4

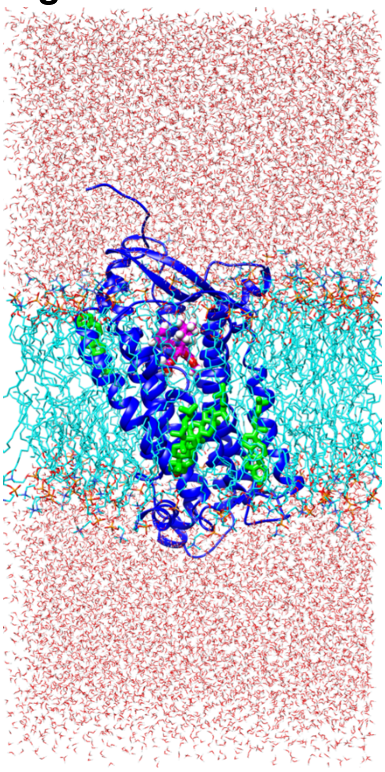

**Figure S4:** Generated system. Blue is hETB (residues 80–404). Magenta, green, and cyan are, respectively, bosentan, cholesterols, and POPC lipid molecules. T4 lysozyme, which was linked to the cytoplasmic side of hETB in the X-ray structure (5x93; see figure S3), was removed. Values of three RCs of this conformation are: $\lambda^{(\alpha)} = 8.22$ Å, $\lambda^{(\beta)} = 8.35$ Å, and $\lambda^{(\gamma)} = 6.21$ Å. History of generation of this conformation is given in figure S2, where this conformation is named "Complex structure of figure S4 (native-complex structure)".

#### Figure S5

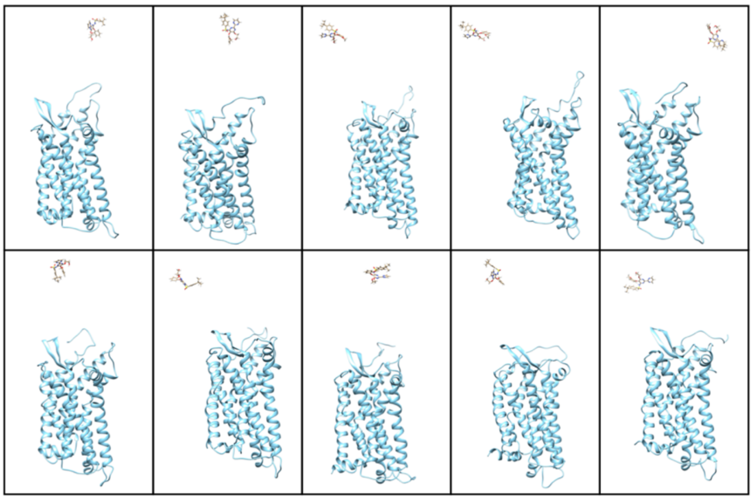

**Figure S5:** Ten conformations from the separation simulations. Only bosentan and hETB are shown removing the other portions. These conformations are called "Unbound conformations in figure S5" in figure S2.

#### Figure S6

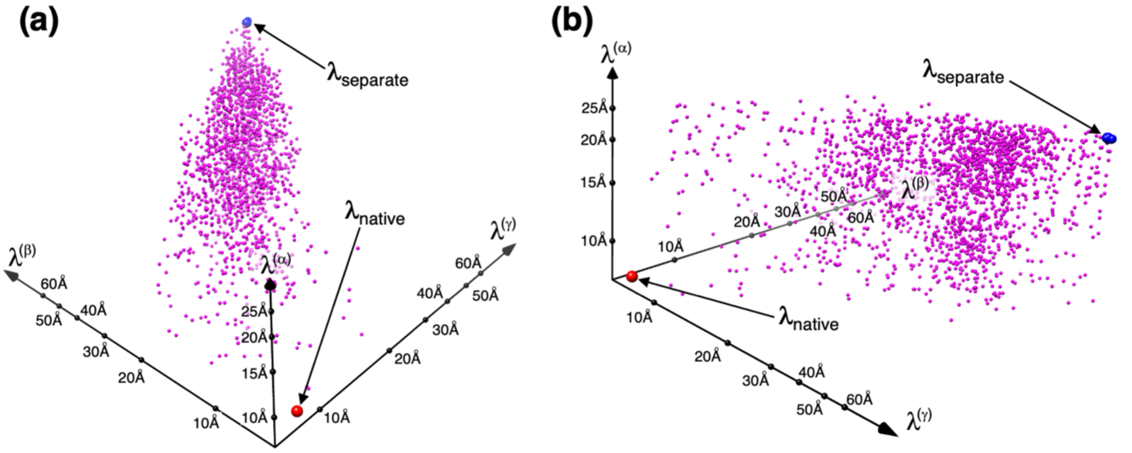


**Figure S6:** Distributions of 2,000 conformations (magenta-colored small spheres) from diffusion simulations. These conformations are named as "Distributed conformations in figure S6" in figure S2. Panels (**a**) and (**b**) are the same distribution viewed from different orientations in 3D-RC space. Blue-colored small spheres labeled "$\lambda_{\text{sepatate}}$" are positions of ten dissociated conformations from separation simulations, which are the initial conformations of diffusion simulations. Although $\lambda_{\text{sepatate}}$ involves 10 conformations, which are different to one another in the real space as shown in figure S5, those are closed to each other in the 3D-RC space. Red-colored small sphere labeled "$\lambda_{\text{native}}$" is position of native complex structure (figure S4): $\lambda^{(\alpha)}$ = 8.22 Å, $\lambda^{(\beta)}$ = 8.35 Å, and $\lambda^{(\gamma)}$ = 6.21 Å.

#### Table S1. Zone Setting

| zone No. [a)] | zones [b)] | | | | | |
|---|---|---|---|---|---|---|
| $k^{(h)}$ | $[\lambda^{(\alpha)}_{k(\alpha)}]_{min}$ | $[\lambda^{(\alpha)}_{k(\alpha)}]_{max}$ | $[\lambda^{(\beta)}_{k(\beta)}]_{min}$ | $[\lambda^{(\beta)}_{k(\beta)}]_{max}$ | $[\lambda^{(\gamma)}_{k(\gamma)}]_{min}$ | $[\lambda^{(\gamma)}_{k(\gamma)}]_{max}$ |
| 1 | 7.0000 | 8.0000 | 7.0000 | 7.6000 | 6.0000 | 6.6000 |
| 2 | 7.5000 | 8.5500 | 7.3000 | 7.9450 | 6.3000 | 6.9450 |
| 3 | 8.0000 | 9.1000 | 7.6000 | 8.2900 | 6.6000 | 7.2900 |
| 4 | 8.5500 | 9.7050 | 7.9450 | 8.6867 | 6.9450 | 7.6867 |
| 5 | 9.1000 | 10.3100 | 8.2900 | 9.0835 | 7.2900 | 8.0835 |
| 6 | 9.7050 | 10.9755 | 8.6867 | 9.5398 | 7.6867 | 8.5398 |
| 7 | 10.3100 | 11.6410 | 9.0835 | 9.9960 | 8.0835 | 8.9960 |
| 8 | 10.9755 | 12.3730 | 9.5398 | 10.5207 | 8.5398 | 9.5207 |
| 9 | 11.6410 | 13.1051 | 9.9960 | 11.0454 | 8.9960 | 10.0454 |
| 10 | 12.3730 | 13.9104 | 10.5207 | 11.6488 | 9.5207 | 10.6488 |
| 11 | 1.31051 | 1.47156 | 11.0454 | 12.2522 | 10.0454 | 11.2522 |
| 12 | 13.9104 | 15.6014 | 11.6488 | 12.9462 | 10.6488 | 11.9462 |
| 13 | 14.7156 | 16.4872 | 12.2522 | 13.6401 | 11.2522 | 12.6401 |

| | | | | | | |
|---|---|---|---|---|---|---|
| 14 | 15.6014 | 17.4615 | 12.9462 | 14.4381 | 11.9462 | 13.4381 |
| 15 | 16.4872 | 18.4359 | 13.6401 | 15.2361 | 12.6401 | 14.2361 |
| 16 | 17.4615 | 19.5077 | 14.4381 | 16.1538 | 13.4381 | 15.1538 |
| 17 | 18.4359 | 20.5795 | 15.2361 | 17.0715 | 14.2361 | 16.0715 |
| 18 | 19.5077 | 21.7585 | 16.1538 | 18.1269 | 15.1538 | 17.1269 |
| 19 | 20.5795 | 22.9374 | 17.0715 | 19.1822 | 16.0715 | 18.1822 |
| 20 | 21.7585 | 24.2343 | 18.1269 | 20.3959 | 17.1269 | 19.3959 |
| 21 | 22.9374 | 25.5312 | 19.1822 | 21.6096 | 18.1822 | 20.6096 |
| 22 | | | 20.3959 | 23.0053 | 19.3959 | 22.0053 |
| 23 | | | 21.6096 | 24.4010 | 20.6096 | 23.4010 |
| 24 | | | 23.0053 | 26.0061 | 22.0053 | 25.0061 |
| 25 | | | 24.4010 | 27.6112 | 23.4010 | 26.6112 |
| 26 | | | 26.0061 | 29.4570 | 25.0061 | 28.4570 |
| 27 | | | 27.6112 | 31.3028 | 26.6112 | 30.3028 |
| 28 | | | 29.4570 | 33.4255 | 28.4570 | 32.4255 |
| 29 | | | 31.3028 | 35.5482 | 30.3028 | 34.5482 |
| 30 | | | 33.4255 | 37.9894 | 32.4255 | 36.9894 |
| 31 | | | 35.5482 | 40.4305 | 34.5482 | 39.4305 |
| 32 | | | 37.9894 | 43.2378 | 36.9894 | 42.2378 |
| 33 | | | 40.4305 | 46.0451 | 39.4305 | 45.0451 |
| 34 | | | 43.2378 | 49.2734 | 42.2378 | 48.2734 |
| 35 | | | 46.0451 | 52.5018 | 45.0451 | 51.5018 |
| 36 | | | 49.2734 | 56.2145 | 48.2734 | 55.2145 |
| 37 | | | 52.5018 | 59.9271 | 51.5018 | 58.9271 |
| 38 | | | 56.2145 | 64.1966 | 55.2145 | 63.1966 |
| 39 | | | 59.9271 | 68.4661 | 58.9271 | 67.4661 |

a) $k^{(h)}$ ($h = \alpha, \beta, \gamma$; $k^{(h)} = 1, \ldots, n_{vs}(\lambda^{(h)})$) is index to specify zone position along $\lambda^{(h)}$. Numbers of zones for $\lambda^{(\alpha)}$, $\lambda^{(\beta)}$, and $\lambda^{(\gamma)}$ axes are: $n_{vs}(\alpha) = 21$, $n_{vs}(\beta) = 39$, and $n_{vs}(\gamma) = 39$.

b) The lower and upper boundaries for the $k$-th zone along the axis $\lambda^{(h)}$ are denoted as $[\lambda_k^{(h)}]_{min}$ and $[\lambda_k^{(h)}]_{max}$, respectively. Unit of zones is Å. The entire sampling range for the axis $\lambda^{(h)}$ is from $[\lambda_1^{(h)}]_{min}$ to $[\lambda_{n_{vs}(h)}^{(h)}]_{max}$, whose widths, $[\zeta_{n_{vs}(h)}^{(h)}]_{max} - [\zeta_1^{(h)}]_{min}$, are 25.5 Å for $h = \alpha$, 61.5 Å for $h = \beta$, and 61.5 Å for $h = \gamma$.

### Section 3. Some physical quantities

#### Subsection 3.1. Unified coordinate system and spatial density of bosentan around hETB

The present GA-mD-VcMD simulation produced $114 \times 10^4$ snapshots as mentioned in the main text. To analyze the molecular structure and molecular motions from the snapshots, a unified coordinate system should be defined. Because the trans-membrane helix (TMH) regions of hETB were structurally conserved well during the simulation, we superimposed the TMH regions (figure S7**a**) of each snapshot on those of the native complex structure (figure S4). The other portions of the system (regions other than the TMH regions of hETB, bosentan, cholesterols, POPCs, water molecules and ions) in the snapshots were moved according to this superposition. This procedure corresponds to define a unified coordinate system on the TMH regions (a body-fixed coordinate system). This superposition was applied to all the snapshots, and physical quantities were calculated using those superposed snapshots.

Benefit of GA-mD-VcMD is that an equilibrated thermodynamic weight (statistical weight) at a simulation temperature (300 K in the present study) is assigned to each snapshot (equation 31 of Ref. 7). Here, we denote the thermodynamic weight assigned to snapshot $i$ as $w_i$. The position of the bosentan's centroid ($CMb$) is referred to as $\boldsymbol{r}_{CMb}$, and $\boldsymbol{r}_{CMb}$ of snapshot $i$ as $\boldsymbol{r}_{CMb,i}$. A method to calculate a spatial density of $\boldsymbol{r}_{CMb}$ around hETB is given as follows: First, the 3D real space is divided into cubes whose volume $\Delta V$ is $3\ \text{Å} \times 3\ \text{Å} \times 3\ \text{Å}$. The position of a cube is represented by its cube–center $\boldsymbol{r}_{cube}$ in 3D real space. Then, the snapshot $i$ is assigned to a cube that involves $\boldsymbol{r}_{CMb,i}$. Assigning all the snapshots to cubes, we calculate the spatial density $\rho_{CMb}(\boldsymbol{r}_{cube})$ of the bosentan's centroid as:

$$\rho_{CMb}(\boldsymbol{r}_{cube}) = \sum_i w_i \delta_{CMb}(\boldsymbol{r}_{cube}; i) \ . \tag{S1}$$

The function $\delta_{CMb}(\boldsymbol{r}_{cube}; i)$ is a delta function defined as:

$$\delta_{CMb}(\boldsymbol{r}_{cube}; i) = \begin{cases} 1 & (\text{if } \boldsymbol{r}_{CMb,i} \text{ is in cube } \boldsymbol{r}_{cube}) \\ 0 & (\text{else}) \end{cases} \ . \tag{S2}$$

Last, we normalized $\rho_{CMb}(\boldsymbol{r}_{cube})$ so that the maximum density is 1: $\rho_{CMb}(\boldsymbol{r}_{cube})|_{max} = 1$. The resultant $\rho_{CMb}(\boldsymbol{r}_{cube})$ is the density assigned to the cube $\boldsymbol{r}_{cube}$.

Also, we define the spatial density of the centroid of a molecular portion $X$ other than the whole of bosentan. Denoting the centroid of $X$ of snapshot $i$ as $\boldsymbol{r}_{CMX,i}$, the spatial density of $X$ is defined as:

$$\rho_{CMX}(\boldsymbol{r}_{cube}) = \sum_i w_i \delta_{CMX}(\boldsymbol{r}_{cube}; i) \ , \quad \text{(S3)}$$

where $\delta_{CMX}$ is a delta function defined as

$$\delta_{CMX}(\boldsymbol{r}_{cube}; i) = \begin{cases} 1 & (\text{if } \boldsymbol{r}_{CMX,i} \text{ is in cube } \boldsymbol{r}_{cube}) \\ 0 & (\text{else}) \end{cases} . \quad \text{(S4)}$$

For instance, in this study, $X$ is set to the tip (Ser 80) of the N-terminal tail of hETB ($X = Nt$) or the tip (Lys 170) of the $\beta$-hairpin of hETB ($X = \beta h$). Then, $\rho_{CMX}(\boldsymbol{r}_{cube})$ is denoted as $\rho_{CMNt}(\boldsymbol{r}_{cube})$ and $\rho_{CM\beta h}(\boldsymbol{r}_{cube})$, respectively.

#### Subsection 3.2. Distance distribution function and radial distribution function

Here, we present a method to calculate a distance distribution function $P_{DDF}(r)$ and radial distribution function $P_{RDF}(r)$, where $r$ is a variable to define the functions. In this study, $r$ is the root-mean-square-deviation $RMSD_{all}^{heavy}$ or $RMSD_{core}^{heavy}$ of bosentan between a snapshot and the native complex structure. $RMSD$ is defined as: $RMSD_{\alpha}^{heavy} = [{N_{atom}}^{-1} \sum_i^{\alpha} (\boldsymbol{r}_i - \boldsymbol{r}_i^{ref})^2]^{1/2}$, where $\boldsymbol{r}_i$ and $\boldsymbol{r}_i^{ref}$ are the $i$-th heavy atom of bosentan in a snapshot and the native complex structure, respectively. For $\alpha = all$, the summation is taken over all heavy atoms in bosentan and $N_{atom}$ is the number of the heavy atoms. For $\alpha = core$, the summation is taken over those in the bosemtan's core region (see figure 3**a** in the main text for the definition of the core region), and $N_{atom}$ is the number of heavy atoms in the core region.

The quantity $P_{DDF}(r)dr$ is a probability to detect the system in a distance window $[r, r + dr]$. The radial distribution function is defined as: $P_{RDF}(r) = P_{DDF}(r)/4\pi r^2$. In this study, we normalized the distribution functions so that the maximum of $P_{DDF}(r)$ or $P_{DDF}(r)$ is set to unity: $[P_{DDF}(r)]_{max} = 1.0$ and $[P_{RDF}(r)]_{max} = 1.0$.

We note that the trans-membrane helices of the snapshot are superposed on those of the native complex structure in advance (see Subsection 3.1 of SI and figure S7**a**) and that no further superposition is applied to the coordinates of the snapshot to calculate $RMSD$. Therefore, $RMSD$ is contributed by three components: Translation of bosentan's centroid (or centroid of the core region) from the snapshot to the native complex, rotation around the bosentan's centroid between the snapshot and the native complex after the translation, and the conformational difference of bosentan between the snapshot and the native complex after the rotation. We note that $RMSD$ converges to the translation with bosentan with bosentan going away from the native-complex position, and that the translation is regarded as a distance between two centroids between the snapshot and the native complex, which is denoted as $r_{bb}$ in figure S7**c**: $P_{DDF}(RMSD) \to P_{DDF}(r_{bb})$ and $P_{RDF}(RMSD) \to P_{RDF}(r_{bb})$.

#### Subsection 3.3. Bosentan–tail contact ratio

A quantity named "cube-based bosentan–tail contact ratio" is introduced as follows: We calculated the minimum heavy-atomic distance, $r_{b-N}$, from bosentan to the N-terminal tail (Ace-Ser 80-Pro 81-Pro 82-Arg 83-Thr 84-Ile 85-Ser 86-Pro 87-Pro 88-Pro 89; Ace is the acetyl group introduced to cap the N-terminal of the N-terminal tail) for snapshot $i$. If $r_{b-N} \leq 5$ Å, the substantial space between bosentan and the N-terminal tail is equal to or smaller than $1$ Å $(= 5$ Å $- 2$ Å $- 2$ Å$)$ assuming that the radius of a heavy atom is $2$ Å approximately. This space of $1$ Å is smaller than the diameter of a water molecule (3 Å). Thus, we judged that bosentan and the N-terminal tail were contacting substantially in the snapshot. Next, we assigned bosentan of the snapshot to a cube $\boldsymbol{r}_{cube}$ that involved the centroid $\boldsymbol{r}_{CMX,i}$ of bosentan. Then, we defined the cube-based bosentan–tail contact ratio:

$$\bar{c}_{b-N}(\boldsymbol{r}_{cube}) = \frac{\sum_i w_i \, c_i \, \delta_{CMb}(\boldsymbol{r}_{cube};i)}{\sum_i w_i \, \delta_{CMb}(\boldsymbol{r}_{cube};i)} \ , \tag{S5}$$

where

$$c_i = \begin{cases} 1 & (\text{if } r_{b-N} \leq 5.0 \text{ Å in snapshot } i) \\ 0 & (\text{else}) \end{cases} . \tag{S6}$$

The ratio $\bar{c}_{b-N}(\boldsymbol{r}_{cube})$ quantifies the bosentan–tail contact ratio in the cube at $\boldsymbol{r}_{cube}$. The larger the $\bar{c}_{b-N}(\boldsymbol{r}_{cube})$ in the cube $\boldsymbol{r}_{cube}$, the higher the contact ratio in the cube. The maximum of $\bar{c}_{b-N}$ is 1.0 (bosentan contacts always to the N-terminal tail when bosentan is in the cube), and the minimum is 0 (always no contact).

Equations S5 and S6 can be used for assessing the bosentan–membrane contact, where $r_{b-N}$ is replaced by $r_{b-m}$, which is the minimum heavy-atomic distance from bosentan to membrane. The function $c_i$ is 1.0 when $r_{b-m} \leq 5.0$ Å is satisfied. We expressed the "cube-based bosentan–membrane contact ratio" as $\bar{c}_{b-m}(\boldsymbol{r}_{cube})$.

#### Subsection 3.4. Spatial patterns of quantity $\Omega$

Here, we defined a method to calculate spatial patterns of a quantity $\Omega$ defined by the coordinates of the system (i.e., $\Omega$ is a function expressed by the system's coordinates): $\Omega = \Omega(\mathrm{x}_1, \mathrm{y}_1, \mathrm{z}_1, \ldots, \mathrm{x}_{N_{all}}, \mathrm{y}_{N_{all}}, \mathrm{z}_{N_{all}})$, where $\mathrm{x}_k$, $\mathrm{y}_k$, and $\mathrm{z}_k$ are, respectively, the x-, y-, and z-coordinates of atom $k$ in the system, and $N_{all}$ is the number of constituent atoms in the system ($N_{all} = 69{,}062$ in the present system). The quantity $\Omega$ is not necessarily defined by all the coordinates but may be done by some of the coordinates. The spatial density is defined as:

$$\overline{\Omega}(\boldsymbol{r}_{cube}) = \frac{\sum_i w_i \Omega_i \delta(\boldsymbol{r}_{cube}; i)}{\sum_i w_i \delta(\boldsymbol{r}_{cube}; i)} \quad , \tag{S7}$$

where $\Omega_i$ is the value of $\Omega$ of snapshot $i$. If the quantity $\Omega$ is a vector, then $\Omega$ is replaced by $\boldsymbol{\Omega}$ in the above equation.

#### Figure S7

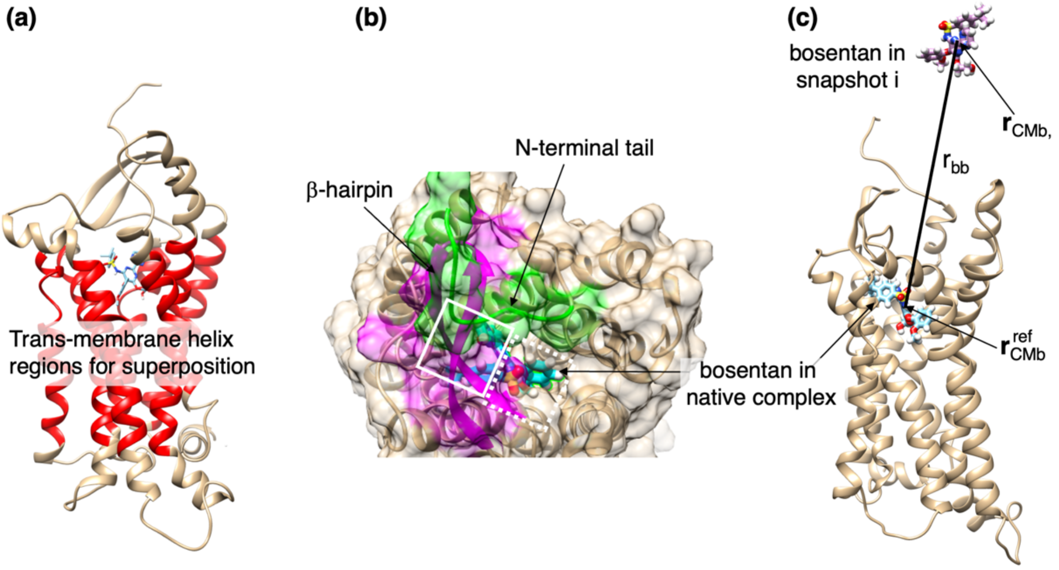


**Figure S7:** Shown structure is native complex (figure S4) for all panels. (**a**) Red-colored segments are trans-membrane helices (TMH) of hETB used for superposition between a snapshot and native complex structure. The superimposed segments are: residues Tyr102–Ile125 in the first TMH (TMH 1), Ile138 –Tyr160 (TMH 2), Lys175–Asp198 (TMH 3), Ala219 –Ile238 (TMH 4), Lys273–Leu295 (TMH 5), Val321–Ser342 (TMH 6), and Leu367–Ala385 (TMH 7). (**b**) Gate of hETB's binding pocket for the native complex structure (figure S4). hETB is presented in ribbon and surface molecular models. N-terminal tail (residues Ser80–Pro89) and β-hairpin (Phe240–His258) are shown in green and magenta, respectively, and the other parts are in ocher for both molecular models. Bosentan in the binding pocket is shown by cyan-colored stick model. In this conformation, a half of the gate of the binding pocket is covered by white solid-line rectangle, where bosentan is not seen directly. In contrast, bosentan is viewed directly in the other half indicated by white broken-line rectangle. (**c**) $r_{bb}$ is distance from bosentan's centroid $\boldsymbol{r}_{CMb,i}$ in snapshot $i$ to that in the native complex structure $\boldsymbol{r}_{CMb}^{ref}$: $r_{bb} = |\boldsymbol{r}_{CMb,i} - \boldsymbol{r}_{CMb}^{ref}|$.

#### Section 4. Distribution of the system's conformation in 3D-RC space

Figure S8 demonstrates the resultant conformational distribution $Q_{cano}(\lambda^{(\alpha)}, \lambda^{(\beta)}, \lambda^{(\gamma)})$ in the 3D-RC space, which was calculated straightforwardly from the simulations. The value of $Q_{cano}$ is provided in a common logarithm. The highest density region (i.e., the lowest free-energy basin) is shown by the red-colored contours in the figure: $\log_{10}[Q_{cano}] = -0.5$, whose free energy is $-RT\ \ln[Q_{cano}] = 0.68$ kcal / mol ($R$ is the gas constant and $T = 300$ K) measured from the lowest free-energy site. Note that the native-complex structure (small black sphere labeled $\lambda_{native}$ in the figure) is located at the periphery of the lowest free-energy basin. The blue-colored region ($\log_{10}[Q_{cano}] = -1.0$; free energy of 1.36 kcal / mol) surrounds the lowest free-energy basin and it is stretched to the direction parallel to the $\lambda^{(\alpha)}$-axis. Remember that $\lambda^{(\alpha)}$ controls the gate width of the hETB's binding pocket (figure 1**b**). Figure S8 indicates that the free-energy slope is gentle to the gate motions. With increasing $\lambda^{(\beta)}$ and $\lambda^{(\gamma)}$, $Q_{cano}$ decreased (i.e., free energy increased).

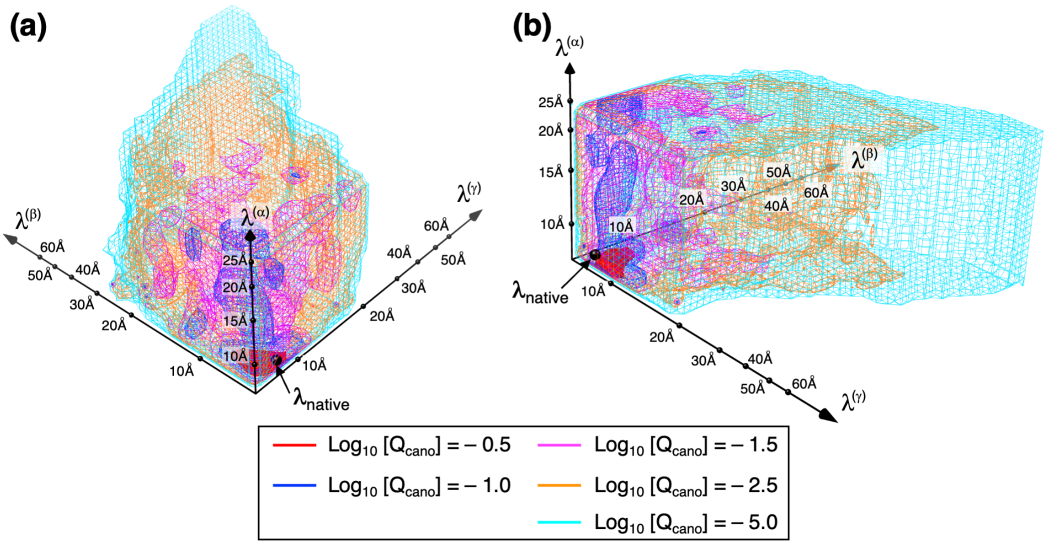


**Figure S8:** Conformational distribution $Q_{cano}(\lambda^{(\alpha)}, \lambda^{(\beta)}, \lambda^{(\gamma)})$ in 3D-RC space obtained from GA-mD-VcMD. The 3D-RC space is constructed by three RCs: $\lambda^{(h)}$ $(h = \alpha, \beta, \gamma)$ (see figure 1 of main text for the definition of RCs). Panels (**a**) and (**b**) are the same distribution viewed from different directions. The distribution is presented in a form of iso-density contour map, and five iso-density levels are displayed in different colors as shown in inset. The density level is given in log scale. Because $Q_{cano}$ is normalized so that the maximum density is set to 1.0, the largest

$\log_{10}[Q_{cano}]$ is zero. Black sphere labeled "$\lambda_{native}$" is position of the native complex structure in the 3D-RC space: $[\lambda^{(\alpha)}, \lambda^{(\beta)}, \lambda^{(\gamma)}] = [8.22\ \text{Å}, 8.35\ \text{Å}, 6.21\ \text{Å}]$. In GA-mD-VcMD, the distribution is defined originally by $Q_{cano}(k^{(\alpha)}, k^{(\beta)}, k^{(\gamma)})$, where $k^{(h)}$ $(h = \alpha, \beta, \gamma)$ is an index to specify the $k^{(h)}$-th zone ranging as $[\lambda^{(h)}_{k^{(h)}}]_{min} < \lambda^{(\alpha)} \leq [\lambda^{(h)}_{k^{(h)}}]_{max}$ (see Table S1). Then, we convert the indices $[k^{(\alpha)}, k^{(\beta)}, k^{(\gamma)}]$ to the RC values as $\lambda^{(h)}_{k^{(h)}} = \{[\lambda^{(h)}_{k^{(h)}}]_{min} + [\lambda^{(h)}_{k^{(h)}}]_{max}\}/2$. The scale of axis (metric) is inhomogeneous because the width of a zone ($[\lambda^{(h)}_{k^{(h)}}]_{max} - [\lambda^{(h)}_{k^{(h)}}]_{min}$) increases with increasing $k^{(h)}$ as shown in Table S1.

#### Section 5. Convergence of distribution

To access the accuracy of the conformational distribution $Q_{cano}(\lambda^{(\alpha)}, \lambda^{(\beta)}, \lambda^{(\gamma)})$ in the 3D-RC space (figure S8), we use a function $E_{local}(\lambda^{(\alpha)}, \lambda^{(\beta)}, \lambda^{(\gamma)})$ (equation 14 of Ref. 7): The smaller the $E_{local}$ at a site $\boldsymbol{\lambda} = [\lambda^{(\alpha)}, \lambda^{(\beta)}, \lambda^{(\gamma)}]$, the more accurate the $Q_{cano}$ in the site and in the vicinity of the site. We judged that a region satisfying $E_{local}(\lambda^{(\alpha)}, \lambda^{(\beta)}, \lambda^{(\gamma)}) < 0.25$ has an appropriate accuracy, where $0.25$ is the criteria used in our previous study.[8,9,10] Here, we denote this region as $R_{acc}(0.25)$.

We performed 55 iterations as mentioned in the main text. Figure S9 illustrates the $R_{acc}(0.25)$ region at iterations 1, 5, 10, 20, 40, and 55. This figure indicates that $R_{acc}(0.25)$ increased rapidly in the first ten iterations. In iteration 10 (figure S9**c**), $R_{acc}(0.25)$ involved both the native complex structure ($\boldsymbol{\lambda}_{native}$) and the completely dissociated conformations ($\boldsymbol{\lambda}_{separate}$). Therefore, we may quit the simulation at iteration 10 to obtain a rough and overall feature of $Q_{cano}$. In fact, the overall shape of $R_{acc}(0.25)$ did not vary largely in iterations 10–55. However, we continued the simulation up to iteration 55 to save more conformations for analysis. Figure S9**f** indicates that $R_{acc}(0.25)$ overlapped well with the regions of $\log_{10}[Q_{cano}] = -5.0$ of figure S8.

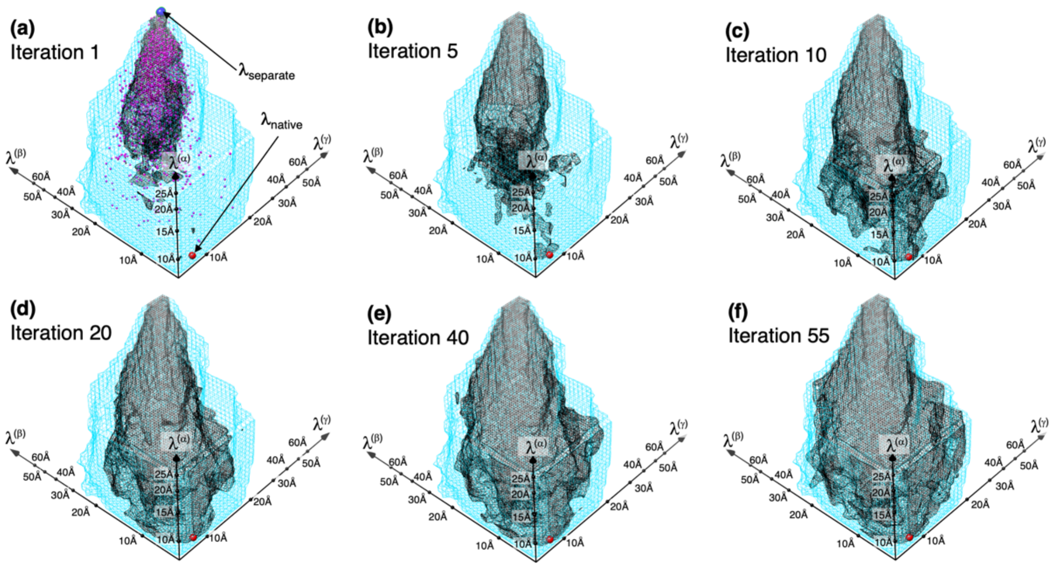

**Figure S9:** Region of $R_{acc}(0.25)$ in six iterations, whose iteration No. are shown near panel labels. See caption of figure 1 of main text for the 3D-RC coordinate axes. Blue-colored small sphere (labeled "$\lambda_{separate}$") in 3D-RC space is completely dissociated conformations from separation simulation (see figures S5 and S6). Red-colored small sphere (labeled "$\lambda_{native}$") in all panels is position of the native-complex structure. Magenta-colored small spheres in panel **a** represent positions of 2,000 conformations from diffusion simulations, which are also shown in figure S6. Cyan-colored iso-density map represents region of $\log_{10}[Q_{cano}] = -5.0$: See figure S8.

#### Section 6. Comparison of the current and previous distributions

We compared the conformational distribution $Q_{cano}\left(\lambda^{(\alpha)}, \lambda^{(\beta)}, \lambda^{(\gamma)}\right)$ with that obtained from our previous study of the ET1–hETB system[2]. The ligand–receptor approaching/departing was regulated by a single RC, $\lambda_2$, in the previous study, whereas it was done by two RCs, $\lambda^{(\beta)}$ and $\lambda^{(\gamma)}$, in the present study. The gate opening/closing of the hETB's binding pocket was controlled by a single RC in both studies: $\lambda_1$ in the present study and $\lambda^{(\alpha)}$ in the previous study. Thus, the conformational distribution was presented two-dimensionally by a function $Q_{cano}^{2D}(\lambda_1, \lambda_2)$ for the ET1–hETB system, although it was expressed by $Q_{cano}(\lambda^{(\alpha)}, \lambda^{(\beta)}, \lambda^{(\gamma)})$ for the bosentan–hETB system.

For the comparison, we contracted $Q_{cano}(\lambda^{(\alpha)}, \lambda^{(\beta)}, \lambda^{(\gamma)})$ to a 2D form $Q_{cano}^{2D}(\lambda^{(\alpha)}, \lambda^{(\beta)})$ or $Q_{cano}^{2D}(\lambda^{(\alpha)}, \lambda^{(\gamma)})$ simply as:

$$Q_{cano}^{2D}\left(\lambda^{(\alpha)}, \lambda^{(\beta)}\right) = \int Q_{cano}(\lambda^{(\alpha)}, \lambda^{(\beta)}, \lambda^{(\gamma)}) d\lambda^{(\gamma)} \quad \text{(S8)}$$

or

$$Q_{cano}^{2D}\left(\lambda^{(\alpha)}, \lambda^{(\gamma)}\right) = \int Q_{cano}(\lambda^{(\alpha)}, \lambda^{(\beta)}, \lambda^{(\gamma)}) d\lambda^{(\beta)} \ . \quad \text{(S9)}$$

Then, the computed 2D distributions are normalized so that the largest value of the distribution, $[Q_{cano}^{2D}]_{max}$, is set to 1: $[Q_{cano}^{2D}]_{max} = 1$. Then, the 2D distribution can be converted to a potential of mean force as:

$$PMF^{2D}\left(\lambda^{(\alpha)}, \lambda^{(Y)}\right) = -RTln\left[Q_{cano}^{2D}\left(\lambda^{(\alpha)}, \lambda^{(Y)}\right)\right] \ , \quad \text{(S10)}$$

where $Y = \beta$ or $\gamma$. Normalization is applied to $Q_{cano}^{2D}$ so that the lowest $PMF^{2D}$ is 0: $[PMF^{2D}]_{min} = 0$.

Figures S10**a** and S10**b** present $PMF^{2D}\left(\lambda^{(\alpha)}, \lambda^{(\beta)}\right)$ and $PMF^{2D}\left(\lambda^{(\alpha)}, \lambda^{(\gamma)}\right)$, respectively. In the both panels, we found a low free-energy path denoted by an arrow labeled "$p_1$", which was parallel to the $\lambda^{(\alpha)}$-axis. Therefore, the gate opening/closing motions follow the low free-energy path with gentle upward/downward slopes. Arrows labeled "$p_2$" and "$p_3$" represent motions along which bosentan approaches/departs hETB. We note that the free-energy slope along arrow $p_2$ is slightly gentler than that along arrow $p_3$. This seems natural because bosentan can approach/depart the binding pocket of hETB with a less cost when the gate is in an open form. This was also observed in the landscape $Q_{cano}(\lambda_1, \lambda_2)$ in the previous study for the ET1–hETB system (Figure 4A in Ref. 12). However, the scales of the free-energy landscapes in the two studies differed largely: The slope in the current bosentan-hETB landscape was about one-tenth of that in the previous ET1-hETB landscape.

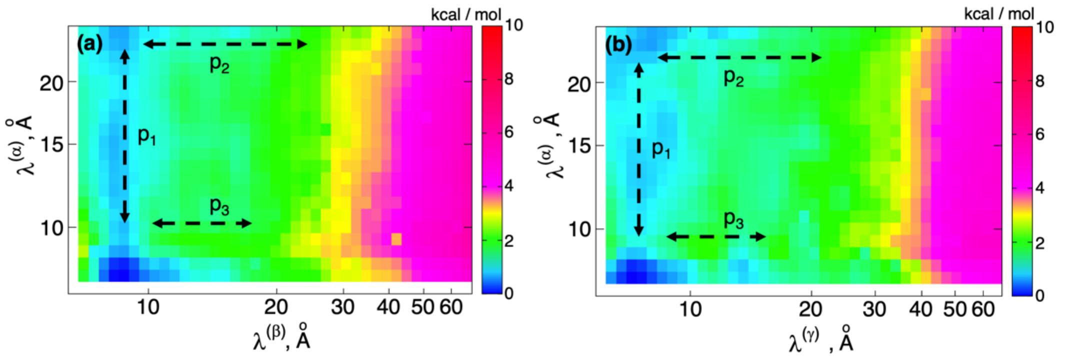


**Figure S10:** 2D distributions (**a**) $Q_{cano}^{2D}\left(\lambda^{(\alpha)}, \lambda^{(\beta)}\right)$ and (**b**) $Q_{cano}^{2D}\left(\lambda^{(\alpha)}, \lambda^{(\gamma)}\right)$ expressed in form of potential of mean force (see Section 5 of SI). Value of $PMF^{2D}$ is presented in color bar. Broken-line arrows with labels $p_1$, $p_2$ and, $p_3$ are called "arrow $p_1$", "arrow $p_2$" and "arrow $p_3$", which are mentioned in main text.

### Figure S11

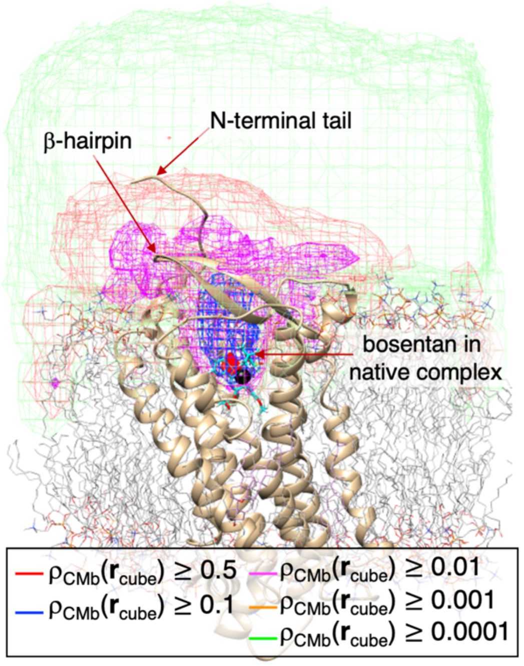


**Figure S11:** Spatial patterns of distribution $\rho_{CMb}(\boldsymbol{r}_{cube})$ of bosentan's centroid $\boldsymbol{r}_{CMb}$ around hETB. Iso-density maps are presented at five contour levels by different colors shown in inset. Shown molecular structure is native complex omitting solvent. Bosentan in the native complex is presented by cyan-colored stick model. N-terminal tail and $\beta$-hairpin are also indicated. See also figure 2 of main text.

#### Figure S12

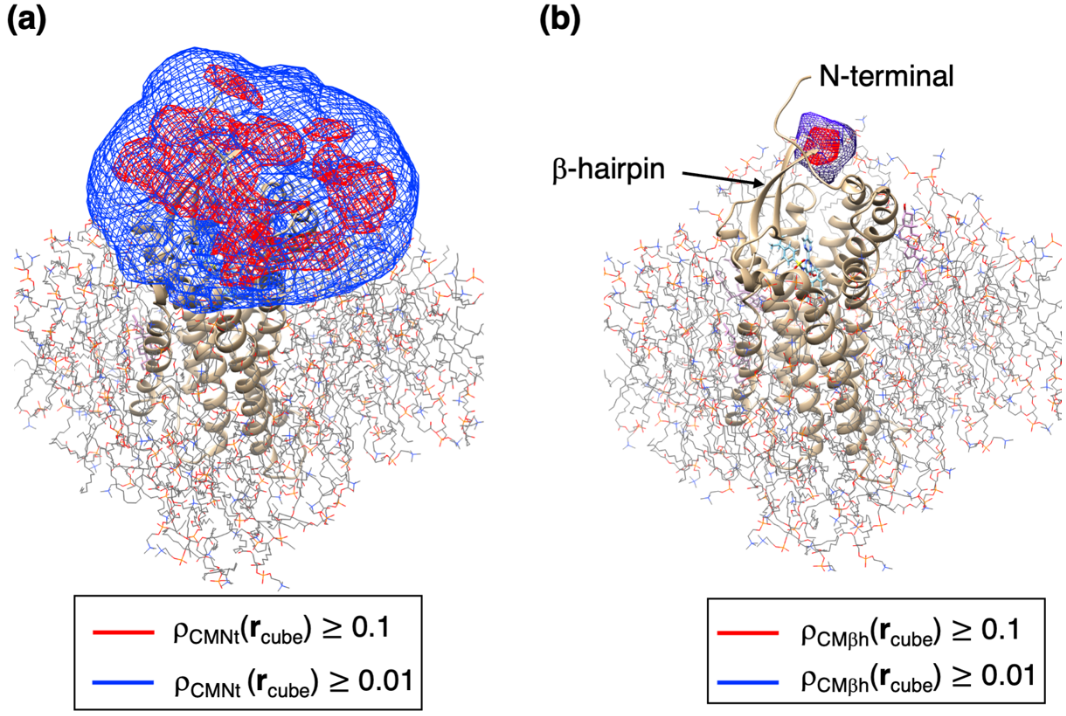


**Figure S12:** (**a**) Spatial density $\rho_{CMNt}(\boldsymbol{r}_{cube})$ for centroid of the tip (Ser 80) of the N-terminal tail and (**b**) $\rho_{CM\beta h}(\boldsymbol{r}_{cube})$ for centroid of the tip (Lys 170) of the $\beta$-hairpin. Contour levels are shown in insets. Shown structure of hETB is the native complex structure (figure S4), and both panels are viewed from the same direction. See Subsection 3.1 of SI for method to calculate $\rho_{CMNt}(\boldsymbol{r}_{cube})$ and $\rho_{CM\beta h}(\boldsymbol{r}_{cube})$.

#### Section 7. Division of $r_{bb}$ axis into slices

We introduced a quantity $r_{bb}$ in figure S7**c**, which is the distance from the bosentan's centroid, $\boldsymbol{r}_{CMb}^{ref}$, in the native-like complex to that, $\boldsymbol{r}_{CMb,i}$, in snapshot $i$. Here, we divide the 3D real space into thirteen slices ($\Delta_k$; $k = 1, \ldots, 13$) along the $r_{bb}$ axis with slice thickness of 5 Å. The actual slice ranges are listed in Table S2. Figure S13 demonstrates distribution of the bosentan's centroid $\boldsymbol{r}_{CMb}$ in each slice.

Apparently, the distribution of the centroid in slices $\Delta_1$,…, $\Delta_4$ are narrow, which indicates that those slices are involved in the binding pocket of hETB. In contrast, the distribution is wide in the slices of $\Delta_5$,…, $\Delta_{13}$. We note that the distribution of centroid varies suddenly at the boundary between $\Delta_4$ and $\Delta_5$, and that this boundary corresponds well to the gate of the binding pocket, which was defined as the boundary between the blue-colored contour region ($\rho_{CMb}(\boldsymbol{r}_{cube}) \geq 0.1$) and the magenta-colored contour region ($\rho_{CMb}(\boldsymbol{r}_{cube}) \geq 0.01$) in figure 2.

#### Table S2. $r_{bb}$-slices $\Delta_k$

| $k$ [a] | $\Delta_k$ | |
|---|---|---|
| | $[r_{bb}^k]_{low}$ [b] | $[r_{bb}^k]_{up}$ [b] |
| 1 | 0 | 5 |
| 2 | 5 | 10 |
| 3 | 10 | 15 |
| 4 | 15 | 20 |
| 5 | 20 | 25 |
| 6 | 25 | 30 |
| 7 | 30 | 35 |
| 8 | 35 | 40 |
| 9 | 40 | 45 |
| 10 | 45 | 50 |
| 11 | 50 | 55 |
| 12 | 55 | 60 |
| 13 | 60 | 65 |

[a] Ordinal number of slices.

[b] $[r_{bb}^k]_{low}$ and $[r_{bb}^k]_{up}$ are lower and upper boundaries for the $k$-th slice $\Delta_k$, respectively, which can be expressed as $[r_{bb}^k]_{low} = 5(k-1)$ and $[r_{bb}^k]_{up} = 5k$. Unit of shown values is Å.

#### Figure S13

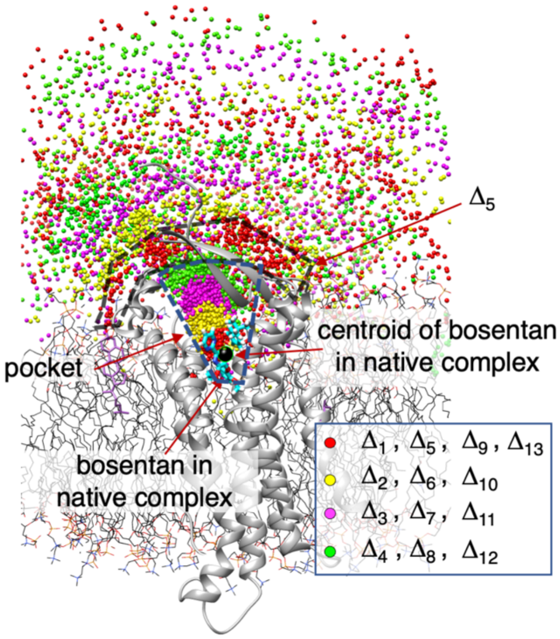


**Figure S13:** Colored dot represent centroids of bosentan in snapshots. Color of a dot corresponds to a $r_{bb}$ slice $\Delta_k$ $(k = 1, \ldots, 13)$, to which the centroid belongs. See Table S2 for actual range of $\Delta_k$. For instance, $\Delta_5$ indicated by broken line is in range of 20 Å $\le r_{bb} <$ 25 Å. Black small sphere represents "centroid of bosentan in native complex" (cyan-colored stick model). Color assigned to $\Delta_k$ is shown in inset, where four colors appear repeatedly as red → yellow → magenta → green with shifting slice as $\Delta_k \rightarrow \Delta_{k+1}$. Five hundred centroids are shown in each slice, which are picked randomly from snapshots in the slice. The shown molecular structure is native complex.

#### Section 8. Conformations picked from some slices

Figure S14 exemplifies snapshots taken from $\Delta_9$ (40 Å $\leq r_{bb} <$ 45 Å), and this slice is involved in the region of $\bar{c}_{b-N} \approx 0.1$ of figure 4. In figures S14**a** and **b**, bosentan contacts to the tip of the N-terminal tail, while it does not in figure S14**c** and **d**. As shown in figure S14**e**, there are snapshots in that bosentan contacts to membrane without touching the N-terminal tail. We also found snapshots in that bosentan contacts to both the N-terminal tail and the membrane (figure not shown). We do not show snapshots in $\Delta_{10} - \Delta_{13}$ because the bosentan–tail contact was formed rarely.

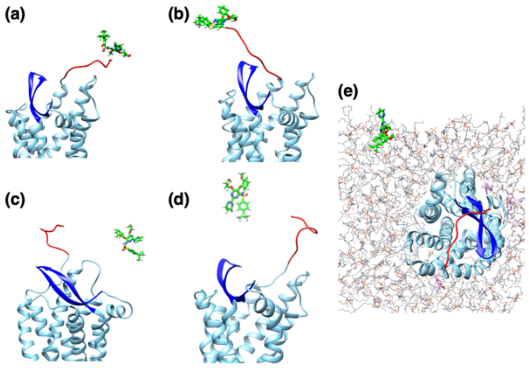


**Figure S14:** Five conformations from $\Delta_9$ (40 Å $\leq r_{bb} <$ 45 Å). Bosentan is distant from hETB's binding pocket. Green-colored stick model represents bosentan. Red- and blue-colored ribbon models are the N-terminal tail and $\beta$-hairpin of hETB, respectively. Solvent is omitted in panels. Panels (**a**)–(**e**) are mentioned in text of SI. Membrane is shown in panel (**e**).

Figure S15 illustrates snapshots in $\Delta_7$ (30 Å $\leq r_{bb} <$ 35 Å), and this range involves the region of $\bar{c}_{b-N} \approx 0.7$. In figures S15**a** and **b**, bosentan contacted to the tail with more area than that in $\Delta_9$ (figures S14**a** and **b**). Therefore, the bosentan–tail contact becomes tightly with bosentan approaching the gate of the pocket. Figure S15**c** illustrates bosentan floating in solvent without contacting to the N-terminal tail, and figure S15**d** does bosentan contacting to both the N-terminal tail and membrane. In figure S15**e**, bosentan buried somewhat in the membrane without contacting to the N-terminal tail.

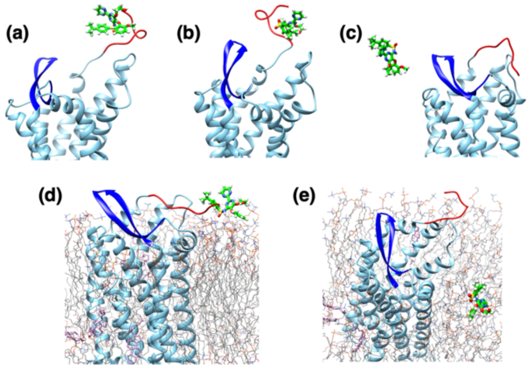


**Figure S15:** Five conformations taken from $\Delta_7$ (30 Å $\leq r_{bb} <$ 35 Å). Bosentan has approached the binding pocket more than those in figure S14. Panels (**a**)–(**e**) are mentioned in text of SI. Membrane is shown in panels (**d**) and (**e**). See also cation of figure S14 for molecular models.

Snapshots in figure S16 are those taken from $\Delta_5$ (20 Å $\leq r_{bb} <$ 25 Å), and this slice involves the region of $\bar{c}_{b-N} \approx 0.9$. This figure exemplifies that bosentan passes the gate with multiple ways. In figures S16**a** and **b**, bosentan is situated in the white solid-line rectangle of figure S7**b** and sandwiched by the N-terminal tail and the $\beta$-hairpin. On the other hand, in figure S16**c** and **d**, bosentan is in the white broken-line rectangle of figure S7**b** when passing the gate. Bosentan in figure S16**c** contacts to the N-terminal tail, although bosentan in figure S16**d** does not. Figure S16**e** is an example that bosentan cannot pass the gate because bosentan is blocked by the $\beta$-hairpin and bosentan should take a roundabout route to reach the gate.

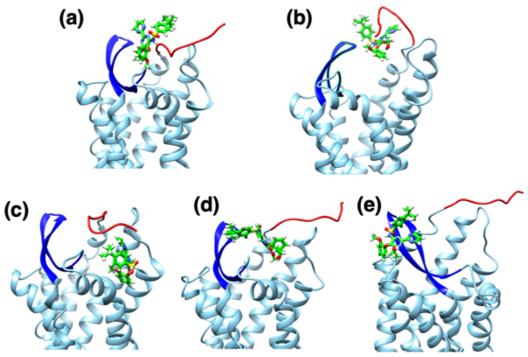


**Figure S16:** Five conformations taken from $\Delta_5$ (20 Å $\leq r_{bb} <$ 25 Å). Bosentan has approached the binding pocket more than those in figure S15. Panels (**a**)–(**e**) are mentioned in text of SI. See also cation of figure S14 for molecular models.

Figure S17 illustrates bosentan's conformations in $\Delta_3$ (10 Å $\leq r_{bb} <$ 15 Å), where bosentan is in the binding pocket with $\bar{c}_{b-N} \approx 0.1$. In Figures S17**a**–**c**, bosentan is between the roots of the N-terminal tail and the $\beta$-hairpin. The N-terminal tail in Figure 10A does not contact to bosentan, although the tail in Figures S17**b** and **c** contacts to bosentan. On the other hand, in figures S17**d** and **e**, bosentan is passing the gate without contacts to either the N-terminal tail or the $\beta$-hairpin. The bosentan–tail contact ratio decreases in the binding pocket as shown in figure S14. This may suggest that that the role of the bosentan–tail contact decreases when bosentan is in the binding pocket.

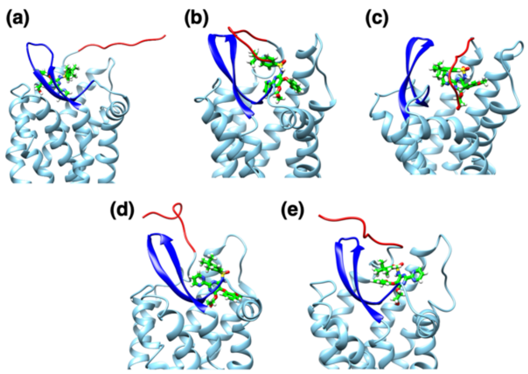


**Figure S17:** Five conformations taken from $\Delta_3$ (10 Å $\leq r_{bb} <$ 15 Å). Bosentan has approached the binding pocket more than those in figure S16. See also cation of figure S14 for molecular models. Panels (**a**)–(**e**) are mentioned in text of SI.

#### Section 9. Residue-based and atom–residue bosentan–tail contact

To analyze the contact of bosentan to a residue of the N-terminal tail as a function of the $r_{bb}$ slice $\Delta_k$, we introduce a "residue–based bosentan–tail contact ratio" $\rho_{cnt}^{\Delta_k}$ as follows: First, the minimum heavy-atomic distance between bosentan and residue $j$ of the tail is calculated for snapshot $m$. Then, if the distance is smaller than or equal to 5 Å, then we judge that the $j$-th residue contacts to bosentan. The residue–based contact ratio is defined as:

$$\rho_{cnt}^{\Delta_k}(j) = \frac{\sum_m w_m c_m(j) D(\Delta_k, m)}{\sum_m w_m} \quad , \tag{S11}$$

where the summations in Eq. S11 are taken over all sampled snapshots. Remember that $w_m$ is the thermodynamic weight assigned to snapshot $m$. $D(k, m)$ is a delta function:

$$D(\Delta_k, m) = \begin{cases} 1 & (\text{if snapshot } m \text{ is involved in } \Delta_k) \\ 0 & (\text{else}) \end{cases} . \tag{S12}$$

The function $c_m(j)$ is defined as:

$$c_m(j) = \begin{cases} 1 & (\text{if residue } j \text{ contacts to bosentan in snapshot } m) \\ 0 & (\text{else}) \end{cases} . \tag{S13}$$

Because the normalization of equation S11 was done by the entire probability (i.e., denominator $\sum_m w_m$ of equation S11 is the probability contributed by all snapshots), we can compare the ratios between different slices: I.e., if $\rho_{cnt}^{\Delta_k}(j) > \rho_{cnt}^{\Delta_{k'}}(j)$, the contact formed at residue $j$ in $\Delta_k$ is stabler than that in $\Delta_{k'}$.

Then, we convert this contact ratio to a potential of mean force ($PMF$) for a free-energy landscape as:

$$F^{\Delta_k}(j) = -RTln\left[\rho_{cnt}^{\Delta_k}(j)\right] \quad . \tag{S14}$$

Then, the inequality $\rho_{cnt}^{\Delta_k}(j) > \rho_{cnt}^{\Delta_{k'}}(j)$ is converted to $F^{\Delta_k}(j) < F^{\Delta_{k'}}(j)$. This quantity is again normalized so that the lowest value of $PMF$ is set to zero in a 2D free-energy landscape constructed by $j$ and $\Delta_k$: $F^{\Delta_k}(j)\big]_{min} = 0$.

Next, to elucidate the bosentan's site specificity contacting to the N-terminal tail, we introduce an "atom–residue contact rate" $\theta_{cnt}^{\Delta_k}(i,j)$, which is the contact ratio of a heavy atom $i$ of bosentan to residue $j$ of the N-terminal tail in the slice $\Delta_k$:

$$\theta_{cnt}^{\Delta_k}(i,j) = \frac{\sum_m w_m c_m(i,j) D(\Delta_k, m)}{\sum_m w_m D(\Delta_k, m)} \quad , \tag{S15}$$

where

$$c_m(i,j) = \begin{cases} 1 & (\text{if residue } j \text{ contacts bosentan's atom } i \text{ in snapshot } m) \\ 0 & (\text{else}) \end{cases} . \tag{S16}$$

Because the normalization in Equation S15 is done over snapshots involved in $\Delta_k$ by the effect of $D(\Delta_k, m)$, the atom–residue contact rate $\theta_{cnt}^{\Delta_k}(i,j)$ should not be compared among different slices. I.e., $\theta_{cnt}^{\Delta_k}(i,j)$ is designed to discuss the bosentan's site specificity in each slice.

#### Figure S18

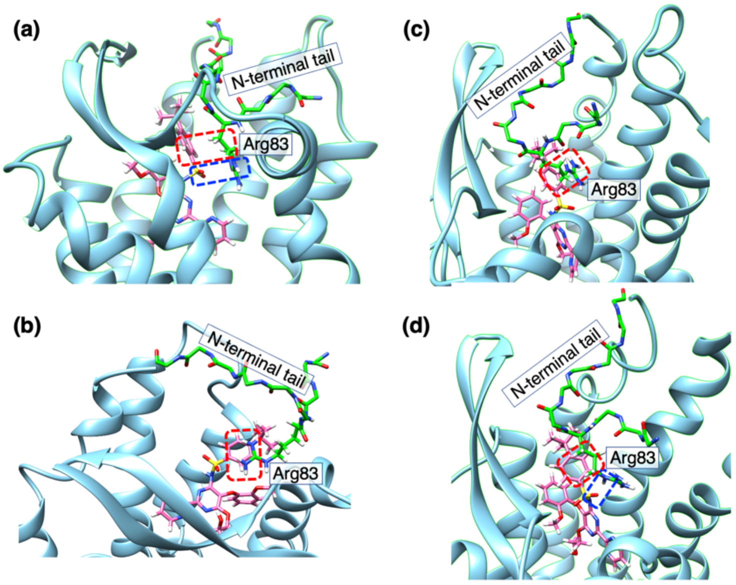


**Figure S18:** (**a**) and (**b**) are snapshots picked from $\Delta_2$, and (**c**) and (**d**) from $\Delta_1$. N-terminal tail and bosentan are presented by green- and magenta-colored stick models, respectively. hETB other than the N-terminal tail is presented by cyan-colored ribbon model. Red-colored broken-line rectangles indicate contacts between hydrophobic sidechain stem of Arg83 in N-terminal tail and hydrophobic atoms 1–10 of bosentan. Blue-colored broken-line rectangles indicate contacts between nitrogen atoms of sidechain tip of Arg83 and oxygen atoms of sulfonamide of bosentan. See figure 6**m** of main text for positions of atoms 1–13 in bosentan.

#### Figure S19

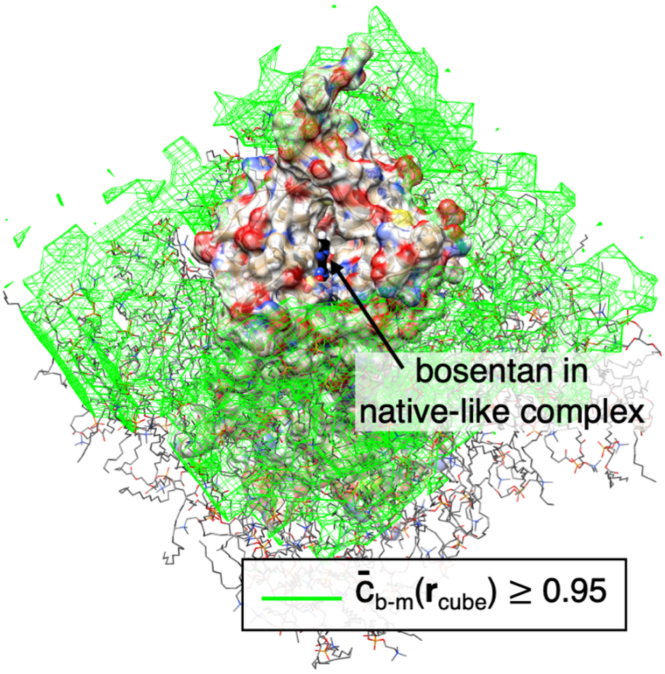


**Figure S19:** Spatial patterns of bosentan–membrane contact ratio $\bar{c}_{b-m}(\boldsymbol{r}_{cube})$: See Subsection 3.3 of SI for definition of $\bar{c}_{b-m}(\boldsymbol{r}_{cube})$. Structure shown is native complex, where bosentan is presented by black stick model, and hETB by surface model. Membrane is presented by stick model. High contact ratio ($\bar{c}_{b-m}(\boldsymbol{r}_{cube}) \geq 0.95$) covers almost entire membrane surface. $\bar{c}_{b-m}(\boldsymbol{r}_{cube})$ decays to zero rapidly with increasing the bosentan–membrane distance $r_{b-m}$ (data not shown).

#### Figure S20

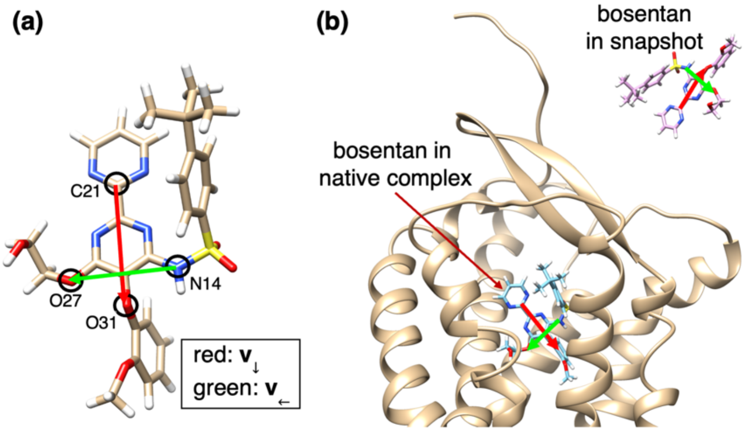


**Figure S20:** (**a**) Two molecular orientation vectors $\boldsymbol{v}_{\downarrow}$ and $\boldsymbol{v}_{\leftarrow}$ fixed on core region of bosentan (figure 3**a**). Two vectors are normalized as $|\boldsymbol{v}_{\downarrow}| = |\boldsymbol{v}_{\leftarrow}| = 1$, and colored as shown in inset. (**b**) Bosentan in native complex and a snapshot. Shown structure of hETB is native complex.

#### Section 10. Inter-molecular native contacts in three X-ray structures.

Figure S21**a** displays the X-ray structures of the bosentan–hETB and bosentan–hETB complexes. The K8794–hETB complex is not shown because it is very similar to the bosentan–hETB complex (figure S3**b**). Bosentan is located at a deeper position of the binding pocket of HETB than the main body (helical part) of ET1 is done (figure S21**a**). This may be an example that the ligands can bind to a deep or shallow position of the binding pocket of a GPCR depending on the ligand type[11]. However, all the three ligands adopt the same binding scheme as discussed in Ref. 1. Figure S21**b** clarifies that the sulphonyl group (SOO) of the sulfonamide of bosentan and K7894 overlap well with the carboxyl group ($COO^{-}$) of the C-terminal residue (Trp 21) of ET1. Besides, the bulky sidechains (ring C defined in figure S21**c**) of bosentan and K7894 overlap well with the sidechain of Trp 21. Therefore, the sulphonyl group and ring C of bosentan play important roles for stabilizing the complex structure.

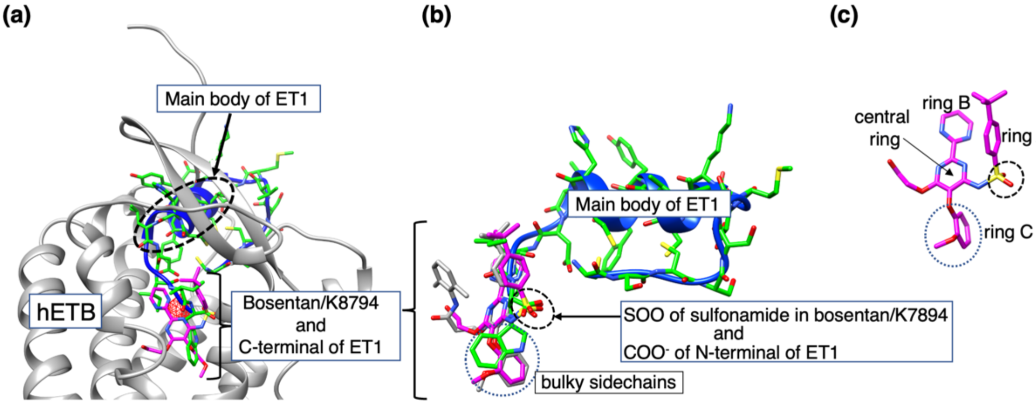


**Figure S21:** Superposition of three complex structures experimentally determined: bosentan–hETB (PDB ID: 5xpr), ET1-hETB (PDB ID: 5glh) and K8794-hETB (PDB ID: 5x93). Superposition is done for the trans-membrane helix regions of hETB (figure S7A), and ligand superposition is not done. (**a**) ET1 is presented by two molecular models: green-colored stick and blue-colored ribbon models. Bosentan is shown by magenta-colored stick model. ET1 is a peptidic ligand consisting of 21 amino-acid residues, and its main body (helix part) is labeled as "Main body of ET1". Positions of bosentan and the C-terminal of ET1, which are overlapped well, are labeled as "Bosentan/K8794 and C-terminal of ET1". K8794 is not shown in this panel, because K8794 and bosentan overlap well (figure S3**b**). hETB is presented by ribbon model in gray. Red-colored contours represent the high-density spot of bosentan's centroid ($\rho_{CMb}(\boldsymbol{r}_{cube}) \geq 0.5$) computed from current simulation. (**b**) Closeup of bosentan, ET1, and K8794 (stick model in gray). Note that three ligands are not superimposed to one another. Instead, as explained in figure S7**a**, the trans-membrane helices of hETB were superimposed to one another, and then hETB was removed. Broken-line circle indicates sulphonyl group ($SOO^{-}$) of sulfonamide of bosentan and K7894, and carboxyl group ($COO^{-}$) of N-terminal of ET1. Dotted-line circle indicates positions of "bulky sidechains" of the three ligands. (**c**) Rings of bosentan are named as "central ring", "ring A", "ring B", and "ring C" as shown in panel. Broken- and dotted-line circles are explained for panel (B). Position of ring C in panel B is at "bulky sidechains".

Figure S22**a** illustrates the X-ray structure of bosentan–hETB focusing on the two oxygen atoms of the sulphonyl group of bosentan, which interact with the sidechain nitrogen atoms of three charged residues of hETB: $N_{\zeta}$ of Lys 182, $N_{\zeta}$ of Lys 273, and $N_{\eta}'s$ of Arg 343. The nitrogen atom ($N14$) of the sulfonamide is also close to $N_{\zeta}$ of Arg 182. We observed similar interaction patterns in the K7894–hETB complex (figure not shown). Figure S22**b**, which focuses on the ET1's C-terminal (Trp 21) of the ET1–hETB complex, demonstrates the interactions between the carboxyl group of Trp 21 and the sidechain nitrogen atoms of Lys 182, Lys 273, and Arg 343. Note that an atom corresponding to N14 of bosentan does not exist in Trp21 of ET1.

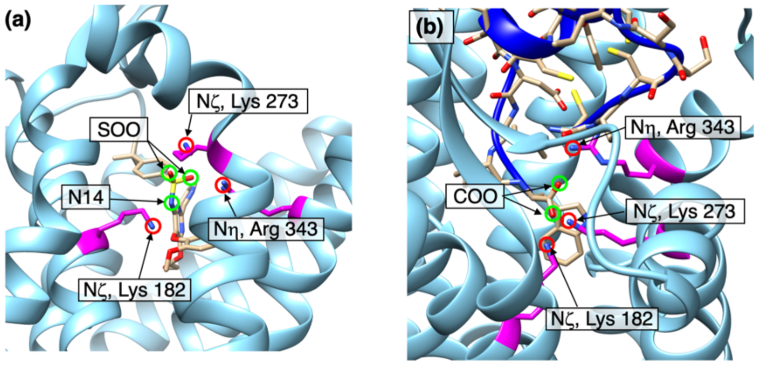


**Figure S22:** (**a**) X-ray structure of bosentan-hETB complex (pdb ID: 5hpr) focusing on bosentan's sulfonamide (SOO), nitrogen (N14), and three charged amino-acid residues, Lys 182, Lys 273, and Arg 343, of hETB (magenta-colored residues). Two nitrogen atoms of arginine ($N_{\eta 1}$ and $N_{\eta 2}$) are named generically as $N_{\eta}$. Green-colored circles indicate two oxygen atoms of sulphonyl group and nitrogen atom of sulfonamide. Red-colored circles do $N\varsigma$ atom of Lys 182, $N\varsigma$ atom of Lys 273, and $N\eta$ atoms of Arg 343. (**b**) X-ray structure of ET1-hETB complex (pdb ID: 5glh) focusing on carboxyl group ($\mathrm{COO^-}$) of C-terminal residue (Trp 21) of ET1. Green- and red-colored circles are oxygen atoms of carboxyl group, which correspond to oxygen atoms in sulphonyl group of bosentan. N14 does not exist in the ET1's C-terminal. Distances among those atoms in the X-ray structures are shown in Table S3. Panels **a** and **b** are drawn from different directions to show clearly the atoms mentioned.

#### Table S3. Atomic distances $r_{\alpha;\beta}$ for four natively contacting contacts in X-ray structures

| | $r_{\alpha;\beta}$ [a)] | | | |
|---|---|---|---|---|
| complex type | $r_{\mathrm{N14;Lys182N_\eta}}$ | $r_{\mathrm{SOO;Lys182N_\zeta}}$ | $r_{\mathrm{SOO;Lys273N_\zeta}}$ | $r_{\mathrm{SOO;Arg343N_\eta}}$ |
| bosentan–hETB | 3.29 Å | 4.56 Å | 3.05 Å | 3.14 Å |
| K8794-hETB | 3.01 Å | 3.41 Å | 2.86 Å | 3.53 Å |
| ET1–hETB | --- | 2.39 Å | 2.24 Å | 2.99 Å |

[a)] $r_{\alpha;\beta}$ is atomic distance between heavy atoms $\alpha$ and $\beta$, which belong to ligand (bosentan, K8794, or ET1) and hETB, respectively: $\alpha = N14$ or SOO of bosentan's sulfonamide as well as $\beta =$ Lys182 $\mathrm{N_\zeta}$, Lys273 $\mathrm{N_\zeta}$, or Arg343 $\mathrm{N_\eta}$. Exact definition for $r_{\alpha;\beta}$ is presented in main text.

#### Section 11. Contacts between heavy atoms $\alpha$ and $\beta$

Given a snapshot $i$, we define a delta function regarding the distance $r_{\alpha;\beta}$ between atoms $\alpha$ and $\beta$:

$$\delta^i_{\alpha;\beta} = \begin{cases} 1 & (\text{if } r_{\alpha;\beta} < 5.0\ \text{Å for snapshot } i) \\ 0 & (\text{else}) \end{cases} . \quad \text{(S17)}$$

The distance of 5.0 Å is a threshold to judge if the atoms $\alpha$ and $\beta$ are contacting (see the main text). Therefore, $\delta^i_{\alpha;\beta}$ is the number of contacts between the two atoms. Next, we introduce another delta function:

$$\delta^i_{r'_{bb}} = \begin{cases} 1 & (\text{if } r_{bb} \text{ of snapshot } i \text{ is in } \mathrm{R}(r'_{bb})) \\ 0 & (\text{else}) \end{cases} , \quad \text{(S18)}$$

where $\mathrm{R}(r'_{bb})$ represents an $r_{bb}$ range of $[r'_{bb} - 0.25\ \text{Å}; r'_{bb} + 0.25\ \text{Å}]$. Then, the expectation value of the number of contacts between atoms $\alpha$ and $\beta$ for snapshots, whose $r_{bb}$ is in $\mathrm{R}(r'_{bb})$, is calculated as:

$$\langle N_{\alpha;\beta}(r'_{bb}) \rangle = \frac{\sum_i w_i \delta^i_{r'_{bb}} \delta^i_{\alpha;\beta}}{\sum_i w_i \delta^i_{r'_{bb}}} . \quad \text{(S19)}$$

$\langle N_{\alpha;\beta}(r'_{bb}) \rangle$ ranges between 0 and 1. If the two atoms always contact to each other for all snapshots in $\mathrm{R}(r'_{bb})$, then $\langle N_{\alpha;\beta}(r'_{bb}) \rangle = 1$. If always no contact, $\langle N_{\alpha;\beta}(r'_{bb}) \rangle = 0$.

As discussed in the main text, we study four distances $r_{\alpha;\beta}$ to analyze the intermolecular interactions between bosentan's sulfonamide and amino-acid residues of hETB in the binding pocket: $r_{\mathrm{N14;Lys182N_\zeta}}$, $r_{\mathrm{SOO;Lys182N_\zeta}}$, $r_{\mathrm{SOO;Lys273N_\zeta}}$, and $r_{\mathrm{SOO;Arg343N_\eta}}$. Then, the expectation values for those four contacts are, respectively, denoted as $\langle N_{\mathrm{N14;Lys182N_\zeta}}(r_{bb}) \rangle$, $\langle N_{\mathrm{SOO;Lys182N_\zeta}}(r_{bb}) \rangle$, $\langle N_{\mathrm{SOO;Lys273N_\zeta}}(r_{bb}) \rangle$, and $\langle N_{\mathrm{SOO;Arg343N_\eta}}(r_{bb}) \rangle$, where $r'_{bb}$ was replaced by $r_{bb}$.

Last, we introduce the expectation value of the number of contacts for the four atom pairs as:

$$\langle N_{cnt}(r_{bb}) \rangle = \frac{\sum_i w_i \delta^i_{r_{bb}} \sum^{pairs}_{\alpha,\beta} \delta^i_{r_{\alpha,\beta}}}{\sum_i w_i \delta^i_{r_{bb}}} , \quad \text{(S20)}$$

where the inner summation (i.e., summation regarding $\alpha$ and $\beta$) is taken over four atom pairs: $(\alpha, \beta) = (\mathrm{N14; Lys182N_\zeta})$, $(\mathrm{SOO; Lys182N_\zeta})$, $(\mathrm{SOO; Lys273N_\zeta})$, and

$(\mathrm{SOO}; \mathrm{Lys343N_\eta})$. The minimum and maximum of $\langle N_{cnt}(r_{bb})\rangle$ are 0 and 4, respectively. The standard deviation of the number of contacts is calculated as:

$$\langle SD_{cnt}(r_{bb})\rangle = \left[\frac{\sum_i w_i \delta^i_{r_{bb}} \left(\sum^{pairs}_{\alpha,\beta} \delta^i_{r_{bb}}\right)^2}{\sum_i w_i \delta^i_{r_{bb}}} - \langle N_{cnt}(r_{bb})\rangle^2\right]^{1/2} - \langle N_{cnt}(r_{bb})\rangle. \quad \text{(S21)}$$

#### Section 12. Formation of each native contact

To investigate the formation of each native contact, we introduced a function $\rho_{bs}(r_{\alpha;\beta}; R(r_{bb}))$, which is the contact probability as a function of $r_{\alpha;\beta}$ using snapshots in a range $R(r_{bb})$. Figure S23 presents six panels of $\rho_{bs}(r_{\alpha;\beta}; R(r_{bb}))$, and the panels are arranged so that bosentan is approaching the native-complex position from the gate of binding pocket: $r_{bb} = 9.25$ Å, 6.25 Å, 3.25 Å, 2.25 Å, 1.25 Å, and 0.25 Å. At $r_{bb} = 9.25$ Å, the probabilities for all the $r_{\alpha;\beta}$ distances were considerably low (figure S23**a**). Decreasing $r_{bb}$, small peaks appeared at $r_{\alpha;\beta} \approx 3$ Å, which indicates formation of the atomic contacts: A peak for $r_{\alpha;\beta} = r_{\mathrm{SOO;Lys273N_\zeta}}$ in $R(6.25\ \text{Å})$ (figure S23**b**), for $r_{\alpha;\beta} = r_{\mathrm{SOO;Arg343N_\eta}}$ in $R(3.25\ \text{Å})$ (figure S23**c**), and for $r_{\alpha;\beta} = r_{\mathrm{SOO;Arg343N_\eta}}$ in $R(2.25\ \text{Å})$ (figure S23**d**). In $R(1.25\ \text{Å})$, the number of peaks increases although their peak heights were still low. Last in $R(1.25\ \text{Å})$, all the four distances inhibited peaks at $r_{\alpha;\beta} \approx 3$ Å or 4 Å, and the peak heights increased (figure 16**e**). This suggests that the complex structure was stabilized by the inter-molecular interactions (i.e., enthalpy) at the bottom of the binding pocket as observed in the X-ray structure.

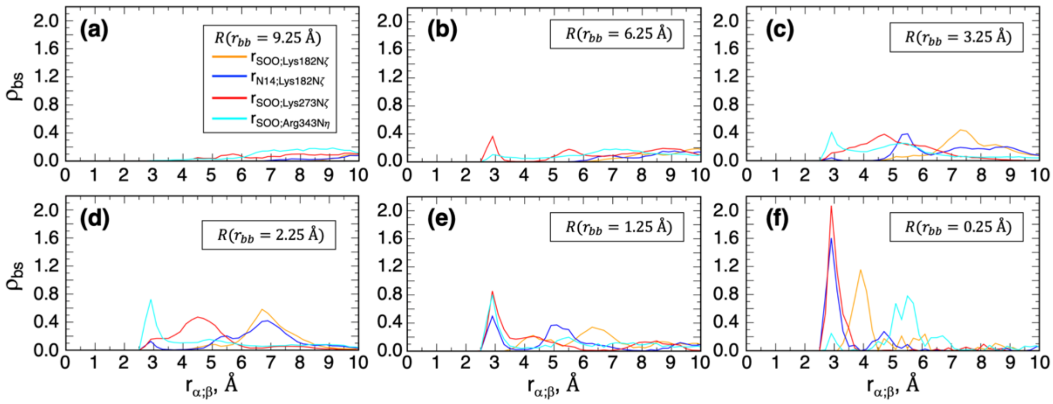


**Figure 23:** Contact probability $\rho_{bs}(r_{\alpha;\beta}; r_{bb})$ as a function of $r_{\alpha;\beta}$ using snapshots in range of (**a**) $R(r_{bb} = 9.25\ \text{Å})$, (**b**) $R(r_{bb} = 6.25\ \text{Å})$, (**c**) $R(r_{bb} = 3.25\ \text{Å})$, (**d**) $R(r_{bb} = 2.25\ \text{Å})$, (**e**) $R(r_{bb} = 1.25\ \text{Å})$, and (**f**) $R(r_{bb} = 0.25\ \text{Å})$. $\rho_{bs}(r_{\alpha;\beta}; r_{bb})$ is normalized as $\int_0^\infty \rho_{bs}(r_{\alpha;\beta}; r_{bb}) dr_{\alpha;\beta} = 1$.

#### Figure S24

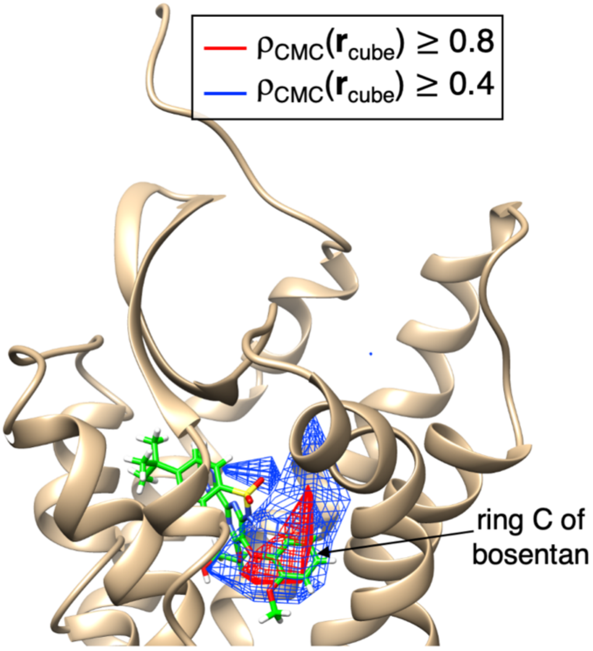


**Figure S24:** Spatial density map for centroid of ring C (see figure S21**c**) of bosentan presented by two contours ($\rho_{CMC}(\boldsymbol{r}_{cube}) \geq 0.8$ and 0.4). Green-colored stick model represents bosentan in native complex.
